# Macroevolutionary diversification rates show time-dependency

**DOI:** 10.1101/396598

**Authors:** L. Francisco Henao Diaz, Luke J. Harmon, Mauro T.C. Sugawara, Eliot T. Miller, Matthew W. Pennell

## Abstract

For centuries, biologists have been captivated by the vast disparity in species richness between different groups of organisms. Variation in diversity is widely attributed to differences between groups in how fast they speciate or go extinct. Such macroevolutionary rates have been estimated for thousands of groups and have been correlated with an incredible variety of organismal traits. Here we analyze a large collection of phylogenetic trees and fossil time series and describe a hidden generality amongst these seemingly idiosyncratic results: speciation and extinction rates follow a scaling law where both depend on the age of the group in which they are measured, with the fastest rates in the youngest clades. Using a series of simulations and sensitivity analyses, we demonstrate that the time-dependency is unlikely to be a result of simple statistical artifacts. As such, this time-scaling is likely a genuine feature of the Tree of Life -- hinting that the dynamics of biodiversity over deep time may be driven, in part, by surprisingly simple and general principles.

**Significance Statement:** Some branches of the Tree of Life are incredibly diverse, while others are represented by only a few living species. Ultimately, this difference reflects the balance of the formation and the extinction of species. Countless explanations have been proposed for why the rates of these two processes vary between lineages including aspects of the organisms themselves and the environments they live in. Here we reveal that a substantial amount of variation in these rates is associated with a simple factor -- time. Younger groups appear to accumulate diversity at much faster rates than older groups. This time-scaling of macroevolutionary rates suggests that there may be hidden generalities governing the diversification of life on Earth.

---

Though once controversial, it is now widely accepted that both the traits of organisms and the environments in which they live can influence the pace of evolution of life on Earth (1–3). In particular, there is tremendous variation between groups of organisms in the rate at which species form and go extinct. This variation is reflected both in the wildly uneven diversity of clades in the fossil record and in the imbalanced shape of the tree of life (2–5). Our estimates of speciation and extinction rates vary by orders of magnitude when comparing different clades, locations, or time intervals.

In turn, researchers have suggested a tremendous array of mechanisms that may have accelerated or slowed the accumulation of biodiversity. These mechanisms include aspects of organisms, such as colour polymorphism (6), body size (7), and many others; the environment, including geographic region (8, 9), temperature (10), and the interactions between the two (11). Taken as a whole, this growing body of work implicates a wide variety of factors that influence speciation and/or extinction rates and suggests that the growth of the Tree of Life has been largely idiosyncratic.

Despite the rapid growth of research on variation in speciation and extinction rates, few synthetic studies have been attempted. As a result, we know little about whether or not there are common factors that predict speciation and extinction rates across diverse taxa (2). One hurdle to synthesis is that studies, especially those using the Tree of Life, often focus more on relative than absolute diversification rates. That is, studies are focused more on whether speciation or extinction rates are higher in one part of a phylogenetic tree than another, rather than attempting to estimate those rates in absolute terms. This makes comparisons across studies difficult or impossible.

Another compelling reason to gather and compare estimates of speciation and extinction rates across clades is the potential for scale-dependence. In all other instances where macroevolutionary rates have been compared, these rates show a pattern of time-dependence, with the fastest rates estimated over the shortest timescales. This time-scale-dependence is apparent in rates of molecular (12) and trait evolution (13, 14) and has even been observed when estimating long-term rates of sedimentation (15). The prevalence of time-scaling of rate estimates in other types of data, along with previous hints that there may be a similar pattern in speciation rates (16–20), suggests that a broader comparison is needed. Here we explore the time scaling of diversification rates using both phylogenetic data from the Tree of Life and paleobiological data from the fossil record and find evidence that there are indeed general scaling laws that govern macroevolutionary dynamics of speciation and extinction.

## Results and Discussion

Using a Bayesian approach that allows for heterogeneity across the phylogeny (21), we estimated speciation and extinction rates across 104 previously published, time-calibrated molecular phylogenies of multicellular organisms which collectively contained 25,864 terminal branches (*SI Appendix*, Table S1). As many other studies have reported, we found substantial variation in both speciation (Fig. 1A) and extinction rates (Fig. S5) across groups ranging from 0.02 to 1.54 speciation events per lineage per million years -- a two-orders of magnitude difference (16, 22–25). Remarkably, a substantial amount of this variation in rates can simply be explained by time alone. We found a strong negative relationship between the mean rates of both speciation and extinction and the age of the most recent common ancestor of a group (regression on a speciation rates: *β* = −0.542; *P* < 0.001; R^2^ = 0.339; extinction rates *β* = −0.548; *P* < 0.001; R^2^ = 0.155; Fig. 2A, B). In general, regardless of the taxonomic identity, ecological characteristics or biogeographic distribution of the group, younger clades appeared to be both speciating and going extinct faster than older groups.

**Figure 1.**
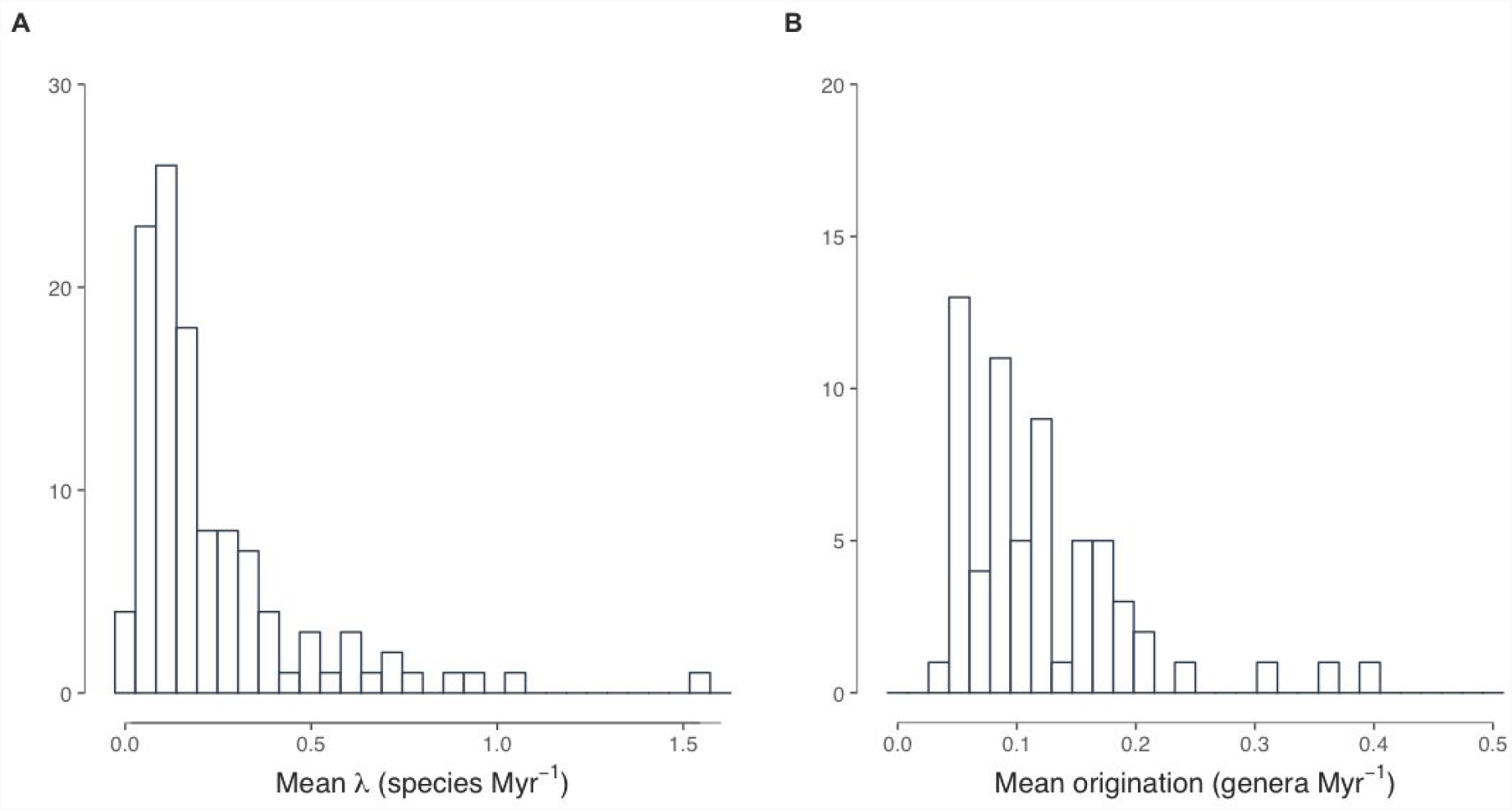
Histogram of mean speciation (λ) and origination rates. A) The lowest speciation rate corresponds to the fern family Matoniaceae (0.02), the highest is Lobelioidae subfamily (1.54). B) The lowest origination rate corresponds to the order Pinales (0.02), the highest is Cingulata order (0.34). Note that axes have different magnitudes.

**Figure 2.**
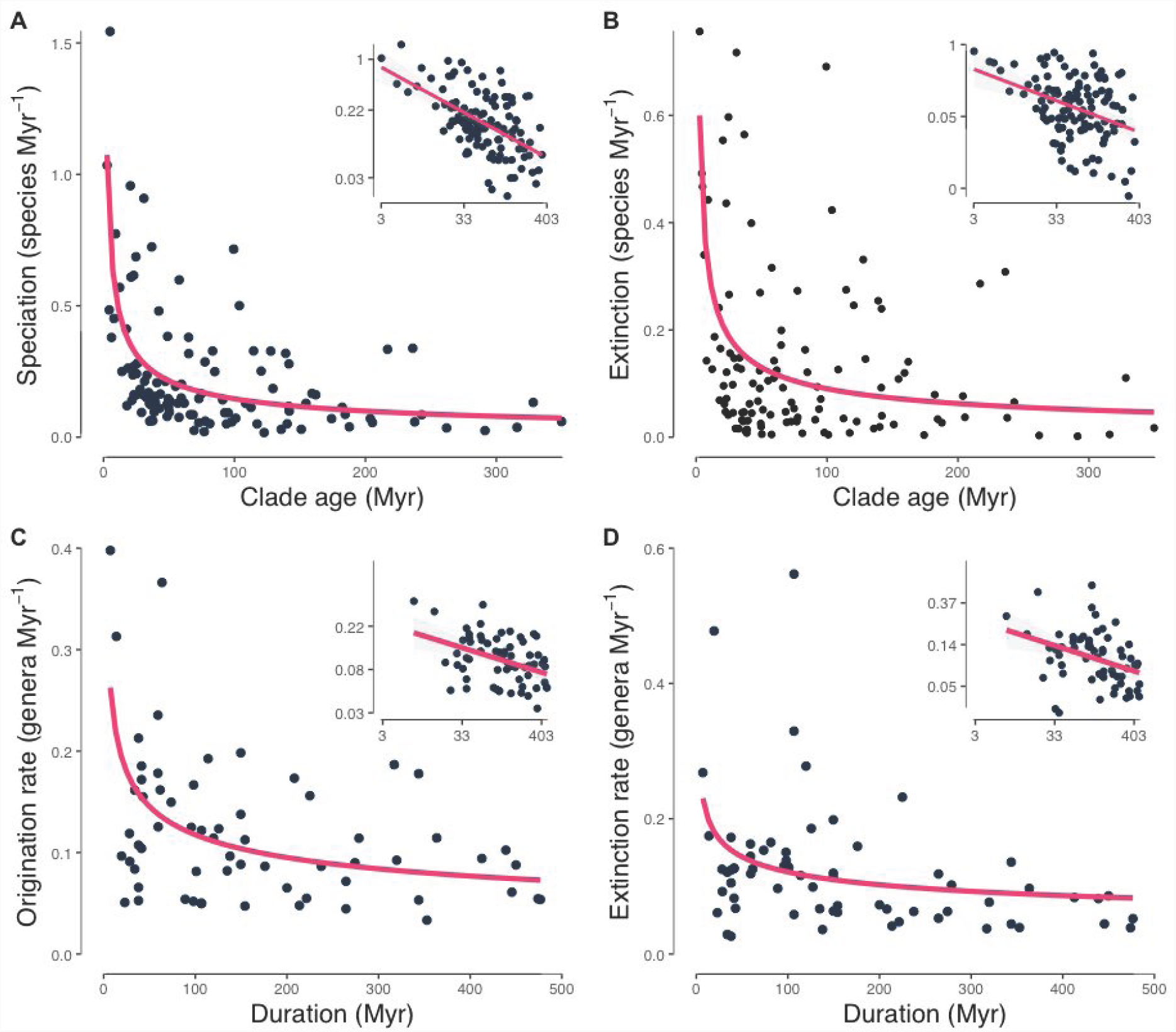
Time-dependency of macroevolutionary rates. Mean per-lineage speciation (Panel A) and extinction (Panel B) rates estimated from 104 molecular phylogenies plotted against the age of the most recent common ancestor of each clade. Panels C and D show the mean rates of origination and extinction of genera estimated from fossil time series from 16 orders of mammals and 24 orders of plants against the total duration the group existed. The insets of each panel show the relationship on a log-log scale.

We also recovered this same scaling pattern using an independent dataset of fossil time series, demonstrating that the time-dependency of diversification rates is a general evolutionary phenomenon and not simply a consequence of using extant-only data. We estimated origination and extinction rates from a curated set of fossil time series consisting of 17 orders of mammals, 22 orders of plants, and 51 orders of marine animals (mostly invertebrates), containing representatives from 6,144 genera (*SI Appendix*, Table S2), using the widely used per capita method (25). For fossil data we measured time as the duration over which a clade of fossil organisms existed and analysed the formation and extinction of genera rather than species. Both origination (*β* = −0.227; *P* < 0.001; R^2^ = 0.152; Fig. 2C) and extinction rates of fossil genera (*β* = −0.245; *P* < 0.01; R^2^ = 0.126; Fig. 2D) were highly dependent on duration.

This result has important consequences for how we measure and interpret rates of diversification. First, it implies that it is not informative to use constant-rate estimators to compare diversification rates of clades of different ages (e.g., (16, 26); this point has been made previously (27, 28) but our results are a particularly clear demonstration that the constant-rate approach is inherently problematic. In recent years, there has been a great deal of progress towards more complex models of diversification which allow for heterogeneity in the process through time and across groups (21, 22, 29). In principle, such variable-process models can alleviate these concerns; if young clades are indeed diversifying rapidly, then the methods will recover this pattern and the analyses will reveal a nested series of upshifts in the rate of diversification (e.g., (30)). In this sense, state-of-the-art methods for measuring diversification rate changes are working exactly as designed. However, our results do call into question how we interpret such results. Typically researchers are not content to merely describe the pattern of rate variation, but rather wish to attribute variation in rates to differences in the organisms or the environments in which they are diversifying. If young groups have faster diversification rates in general, then nearly *any feature* that has recently evolved or *any environment* that organisms have recently colonized will likely be associated with an increase in diversification. It is beyond the scope of the present article to discuss what precisely constitutes evidence of causality in macroevolutionary studies (see Maddison and FitzJohn 2015; Uyeda et al. 2018), but we suggest that it is certainly not a coincidence that many of the regions of the world that are often recognized as hotspots of diversification (at least over the past several million years) are also precisely those regions that harbour young groups of organisms, such as the Páramo of the Andes (31), oceanic islands (32), or polar seas (23).

Potential causes of this ubiquitous pattern fall into two main categories. First, our result may be statistical bias; the underlying process of diversification may be consistent with that assumed by diversification models with constant rates, but our statistical methods are biased towards recovering time-dependence. Second, our result may reflect biological generality; the true dynamics of diversification over macroevolutionary timescales show patterns that are incompletely described by current models and theory and lead to time-dependence in rate estimates. Using a series of simulations and sensitivity analyses, we show that statistical biases are unlikely to account for the entirety of our pattern. We present results investigating a variety of potential artifacts, such as bias in the rate estimators, incomplete sampling, error in divergence time estimation, or acquisition bias (known as the “push of the past” in the macroevolutionary literature), all of which have been invoked when this pattern was previously documented by biologists in studies of individual groups (16–18, 20, 28, 33, 34). While all of these may generate a negative relationship between rate and time, none of the artifactual explanations we considered can fully account for the pattern we find in both phylogenetic and fossil time-series data.

We therefore suggest that the time-dependency of rates is indeed a real phenomenon that requires a biological explanation. Perhaps the simplest, and easiest to dismiss, is that there has been a true, secular increase in rates of diversification through time, such that, globally, macroevolution is faster now than it has ever been in the past. We fit variable rate models to each molecular phylogeny individually but find no evidence of widespread speedups *within* groups (Fig. 3A). While we do find support for shifts in diversification rates within trees (*SI Appendix*, Fig. S1), there is no clear temporal trend. This is inconsistent with the idea of a global speedup. Consistent with previous studies (33), we do not see an increase in rates through time for fossil data (Fig. 3B). Furthermore, previous studies have shown that heterogeneity in rates across groups cannot generate the patterns we observe under realistic diversification scenarios (28).

**Figure 3.**
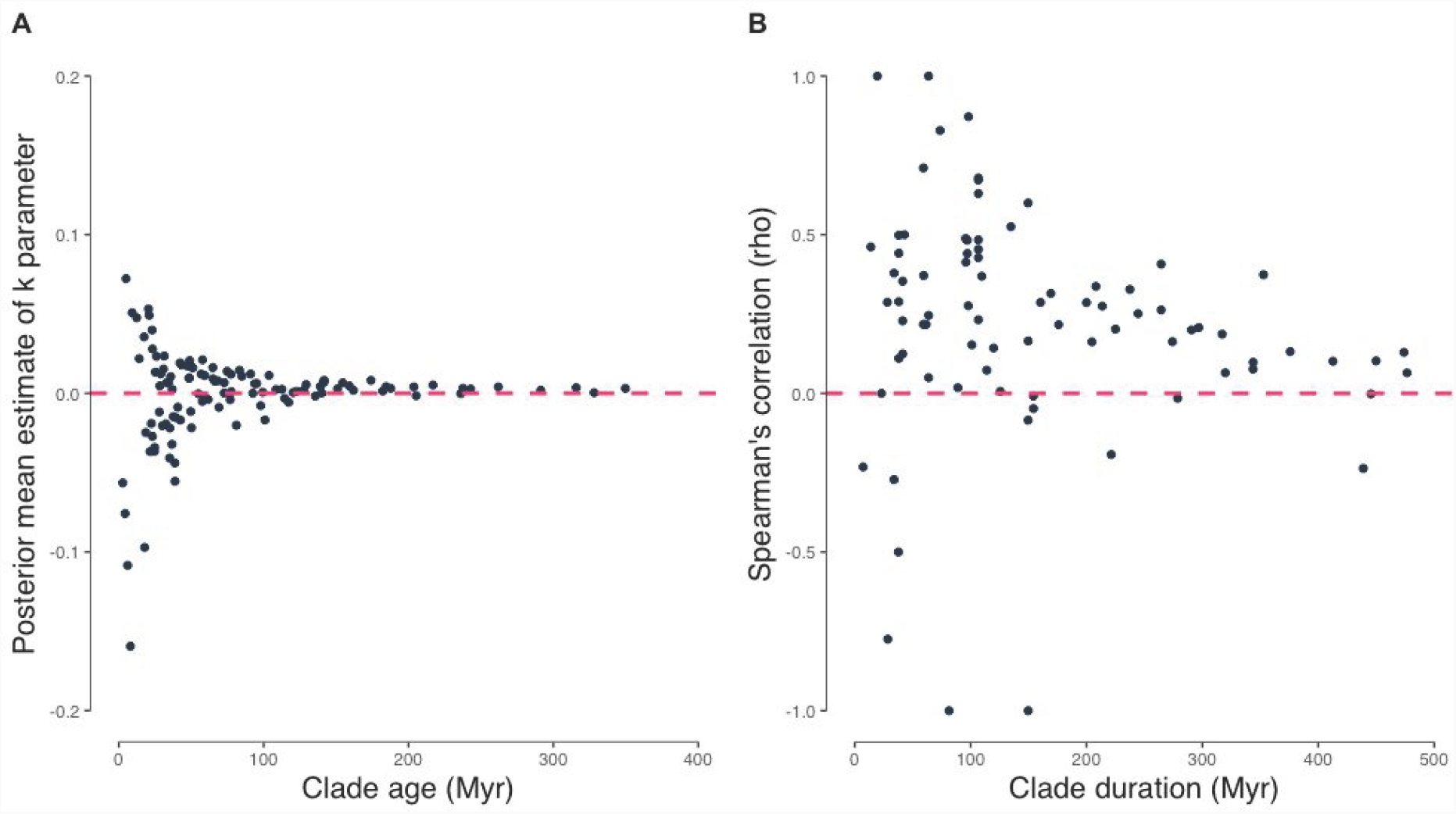
Diversification trends through time. A) Phylogeny mean posterior distribution of shift parameter (k) from BAMM against clade age; mean k value was calculated across each posterior. B) Spearman’s correlation between fossil’s time interval ordination and diversification rate using the per Capita method against clade duration. Neither the k parameter nor Spearman’s correlation coefficients decrease with time. Furthermore, they do not deviate from zero with clade age, suggesting that there is no strong evidence for diversity dependence through time.

The lack of evidence for temporal trends within groups also does not support another commonly invoked explanation for the time-dependency of rates -- that diversity dynamics are shaped by ecological limits (4, 27). If niche or geographic space is constrained, then diversification should slow down as diversity accumulates; this decoupling of clade age and size would then lead directly to time-dependent rates of diversification (28). In this scenario, younger clades are growing near exponentially while older clades have reached stationarity; therefore, the average rates of evolution would thus appear much faster in younger clades relative to older ones (19). This explanation predicts that slowdowns in rates should be ubiquitously observed *within* clades which, as stated above, is incompatible with our findings. Furthermore, the signal for a slowdown should be most apparent in older clades, which is also not apparent in either phylogenetic or paleobiological data (Fig. 3A, B), though we acknowledge that such a signal is particularly difficult to detect in older clades as it may be eroded by subsequent diversification (35). Furthermore, such clade-based explanations depend critically on the premise that higher taxonomic groups (such as families, orders, etc.) are meaningful units for diversification analyses. This may be because taxonomists have actually identified true, evolutionary lineages (36) with dynamics of their own, such as those envisioned by models of taxon cycles (37, 38) or inter-clade competition (39). It might also be the case that taxonomic practice is biased in some yet-unknown way (28). We can evaluate the premise that named clades are special by subsampling our data. Using recently published megaphylogenies of birds (32), ferns (40) and flowering plants (41) we tested whether the slope of the time-dependency of diversification rates was different between named clades and clades descending from randomly chosen nodes of similar age. We found that they were not (Fig. 4), which suggests that clade-specific ecological limits are unlikely to satisfactorily explain our results.

**Figure 4.**
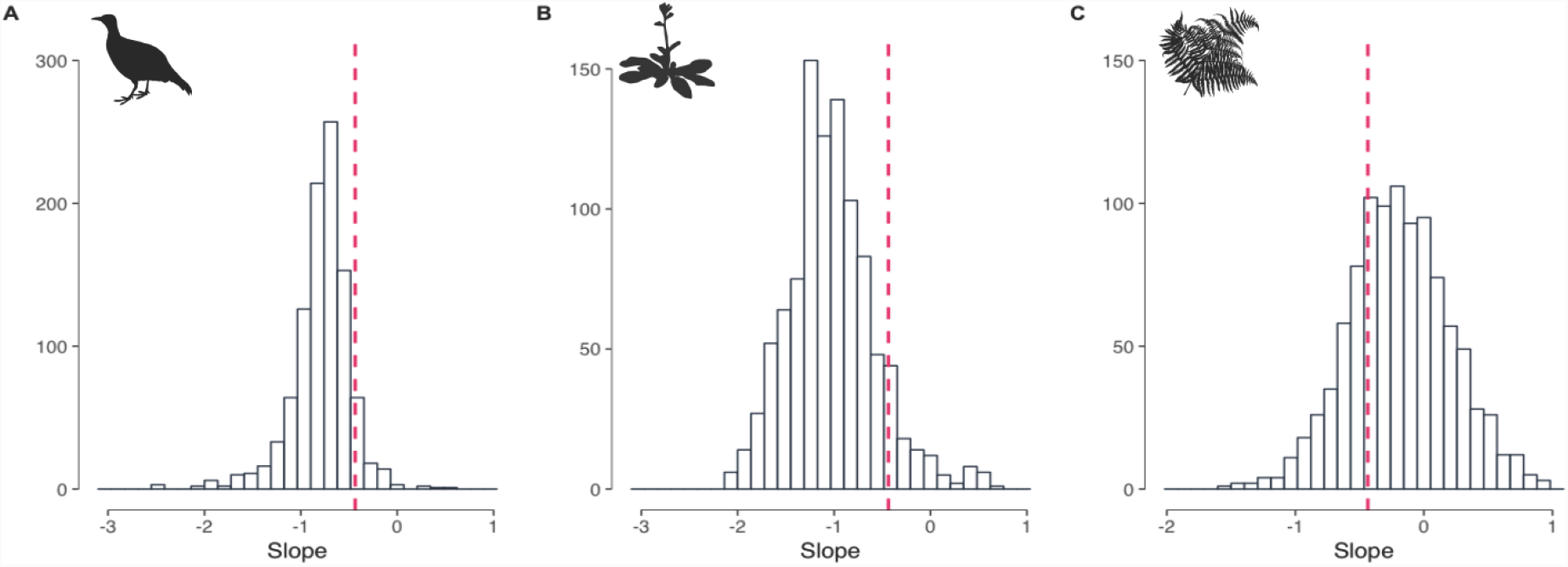
Random clade selection effect. Empirical slope of diversification rate as a function of time, (dashed) compared with the slope’s distribution from randomly sampled clades. In all cases our empirical estimates laid within the negative slopes distribution from randomly drawn clades; showing that our results are not a sampling artifact due to taxonomic delimitation. A) Bird families, B) Angiosperms orders, C) Fern families.

We favor another explanation: that speciation and extinction events are clustered together in time, with these clusters interspersed among long periods where species neither form nor go extinct. For inspiration, we turn to a result, strikingly similar to our own, from an entirely different field. Sadler (15) demonstrated that estimated rates of sediment deposition and erosion are also negatively correlated with the time interval over which they are estimated (this is now known in geology as the “Sadler effect”). This time dependency likely results from the unevenness of sedimentation: geological history is dominated by long hiatuses with no or negative sedimentation (i.e., erosion) punctuated by brief periods where large amounts of sedimentation accumulates (42). Under such a scenario, the mean rate tends to decrease the farther back in time one looks owing to the fact that more and more hiatuses are observed.

We think that Sadler’s rationale could apply to diversification rates as well. There is evidence that extinction events recorded by the fossil record are much more clustered in space and time than we would predict under gradualism (43, 44). Indeed, many of the boundaries of the geological time-scale are defined by large-scale faunal and floral turnover -- the most widely known example of this is undoubtedly the mass extinction event that separates the Cretaceous from the Paleogene. There is abundant evidence, including from our results, that origination and extinction rates tend to be highly correlated over macroevolutionary time, and concentrated in clades with high volatility (33, 45). We expect that if extinction events are concentrated in time, speciation rates will be as well.

We are not the first to propose that pulses of diversification may affect the interpretations of rates estimated on the assumption that diversity accumulates gradually; paleobiologists have found that short geological intervals tend to show higher rates than longer intervals and have argued that this is caused by the concentration of events at the interval boundaries (43, 44). As we noted above, some phylogenetic analyses have uncovered similar patterns (16–20, 34). However, explanations favored by previous work do not explain the entirety of our observation of apparent time-dependency found across both phylogenetic trees and fossils.

Some researchers have suggested that clustered speciation and extinction events observed in the fossil record may be due to large-scale, and possibly regular, climatic fluctuations (46) or an emergent property of complex ecosystems (47). Perhaps a simpler explanation is that species evolve and diversify over a complex geographic landscape, and successful speciation and persistence seem to require the confluence of multiple factors at the right place and time (48). Most speciation events likely occur in lineages with limited or fragmented ranges but the resulting species are also highly prone to go extinct before they can leave their mark on macroevolutionary history (49). This verbal model shares much in common with Futuyma’s model for ephemeral divergence (50), which itself can be invoked to explain the long-observed negative time-dependency of rates of phenotypic evolution (13, 14) and could, potentially, explain a similar pattern in rates of molecular evolution (12). This scenario, consistent with our results, would imply that the scaling of rates of sedimentation, phenotypic divergence, molecular evolution, and diversification with time all might share a common mechanism.

We acknowledge that much more work, both theoretical and empirical, is needed to fully understand the reasons that diversification rates scale with time and this research may or may not support our proposed explanation. In either case, we hope our results call biologists’ attention to surprising regularities in the Tree of Life. Long before well-resolved and robustly dated phylogenies were available for most groups, biologists noted trees had stereotypical features (5, 51) (for a more recent example, see ref. (52)). Macroevolutionary biologists have tended to focus their attention on the particular innovations that may have spurred radiations in particular groups at particular times; a far less explored -- and completely open -- question is why, despite all of the idiosyncrasies and contingencies inherent in the process of diversification, phylogenies from across the Tree of Life look so remarkably similar.

## Methods

### Phylogenetic collection and data cleaning

We collected and curated 104 time-calibrated species-level phylogenies of multicellular organisms from published articles and from data repositories; these trees were fully bifurcating, ultrametric, and contained more than 7 tips and at least 1% of clade’s total richness (*SI Appendix*, Table S1, S3). Trees were checked to ensure branch lengths were in millions of years (53–56). For each group we compiled from the literature the total number of extant species.

### Phylogenetic diversification analysis and regressions

Diversification rates were estimated for each phylogeny using BAMM with default priors and including sampling fraction (57). From the posterior distribution we calculated the mean and variance in speciation and extinction rates across all branches and the frequency of shifts. Model mixing and convergence were assessed by examining the effective sample size in coda (58). In order to calculate the *k* parameter (see *SI Appendix* for more detail), we re-ran BAMM on each phylogenetic dataset and permitted only a single diversification regime per tree. We fit linear models between mean and variance rates on a log-log scale compared with crown ages (estimated from the tree heights). We observed qualitatively identical results when fitting the same linear regressions using the Maximum Likelihood Estimate (MLE) rates for the DR statistic (32).

### Phylogenetic simulations

In order to explore the effect of the ‘push of the past’ (i.e., conditioning on the survival of the clade to the present) we simulated trees with at least 6 taxa and the same ages as our empirical dataset; we used the diversification parameters from the oldest groups (> 150 My), where the curve between rates and age levels off. We repeated this procedure by setting µ = 0.5λ to acknowledge the difficulty in estimating extinction rates and their influence in macroevolutionary dynamics. We repeated both procedures 1000 times.

To evaluate whether the observed time-dependency was a consequence of using named higher taxonomic groups, we develop a novel algorithm for randomly sampling nodes from a tree given the age distribution of a set of named nodes and a tolerance. Using this we were able to compare a temporally equivalent set of unnamed random clades with our empirical tree results. We computed the mean MLE for diversification rates of named clades (across several ranks) from megaphylogenies and to the random clades with equivalent ages. We repeated the latter process 1000 times.

The reconstructed trees usually represent a small proportion of a group’s entire diversity, making diversification analysis sensitive to this sampling fraction. For our purposes, this is especially relevant as older groups tend to be more sparsely sampled than younger ones. Even though previous studies have shown that birth-death estimations are consistent when the sampling fraction is provided (59, 60), we wanted to rule out its effect. We simulated a tree corresponding to each tree in the dataset based on empirical ages, mean rates from phylogenies older that 150 My, and sampling fractions using TreeSim R package (61). We re-estimated the MLE using diversitree (62), repeating this procedure 1000 times.

Errors in ages of young clades can overestimate diversification rates leading to a negative relationship between rates and time. Given the heterogeneous nature of the trees we used, it was not feasible to include uncertainty in branching times (i.e., from a posterior distribution or set of bootstraps). As an alternative, we explored this possibility using simulations; we repeated the simulations as in the ‘push of the past’ section but this time we added an error to clade ages. To each age, we drew a percentage error from a uniform distribution that modified the branch lengths (by addition or subtraction). Simulations were carried with maximum error values from 10% to 90%. In all the simulated cases mentioned we estimated the slope of the log(speciation rate) ∼ log(crown age) regression resulting from each one of the trees; and then we compared these to our empirically estimated slope.

### Fossil collection and cleaning data

We collected and curated fossil information on the first and last appearance of each genus for 39 orders of mammals (17) and plants (22) from online databases. Each order included in our analyses consisted of at least 10 genera, each genus had at least two occurrences, and only occurrences assigned to a unique geological stage were considered (63). We also included 51 orders, each with a minimum of 10 genera, from Sepkoski’s marine animal compendium. In total, our data consists of 6,144 genera distributed across 90 orders that were analysed individually (*SI Appendix*, Table S2).

### Fossil diversification analysis and regressions

Diversification rates were estimated using Foote’s per-capita method (25), which uses the first and last occurrence of the genera to estimate the rates of origination (i.e., rate of appearance of new genera) and extinction for each geological stage. We divided the diversification rate (i.e., origination minus extinction) by the duration of each interval, then estimated the average diversification rate for the orders. We also estimated the correlation between the rate estimates and the time-bin ordination. We fit linear models on log-log scale between average diversification rate and clade duration.

## ACKNOWLEDGMENTS

We thank D. Schluter, S. Magallon, J. Davies, S. Otto, A. Mooers, K. Kaur, B. Neto-Bradley, F. Mazel, J. Rolland, J. Uyeda, and two anonymous reviewers for comments on this work. These analyses were made possible by Compute Canada. This work was funded by a NSERC Discovery Grant to MWP and a NSF Grant DEB #1208912 to LJH.

L. Francisco Henao Diaz ORCID: 0000-0002-1975-490

## Supporting Information

### METHODS

#### Curation of phylogenetic datasets

We collected and curated a set of time-calibrated species level phylogenies from the supplementary materials of published journal articles and from data repositories. Since we needed time-calibrated trees, with branch lengths in standardized units (millions of years), we were unable to rely exclusively on existing compilations (e.g., TreeBase (64), OpenTree (65)). It was important that we gather data from as many different, independent sources as feasible rather than simply slicing up previously constructed “megaphylogenies” (66) for two reasons. First, very large trees are necessarily constructed using only the few genes that are widely available for many taxa; there is thus likely to be substantial uncertainty in both topology and divergence times (66–68). Second, we wanted to minimize the correlation of error across datasets.

Our goal in collecting trees was to assemble a dataset that was as taxonomically unbiased as possible -- so as to avoid focusing on groups that have been previously identified as being adaptive radiations, etc. -- and made up primarily of recent studies under the assumption that, in general, more recently estimated trees tend to be more robust than older ones. To do this, we took two approaches; i) first we systematically surveyed issues of eight journals that routinely publish primary phylogenies (the issues we examined are listed in *SI Appendix*, Table S3); and ii) we searched Open Tree of Life (65), Figshare (figshare.com), and Dryad (datadryad.org) using the following tags: “time-calibrated”, “dated-tree”, and “species-time-tree”. In order to be included, phylogenies had to be fully bifurcating and ultrametric, include more than 7 tips and contain more than 1% of the total diversity of the group being studied.

Through this process we compiled 104 phylogenies from 69 studies, most of them at family (54), genus (19) and order (11) rank; plant (50) and animal (54) groups were nearly evenly represented (*SI Appendix*, Table S1). The number of taxa included in the trees varied from 7 to 4,161 species, and clade (crown) ages range from 2.74 to 349.8 million years (Myr).

For each group represented by a phylogeny, we searched the literature to estimate the total number of extant taxa in the group. Because speciation rates likely have radically different meanings for unicellular organisms (but see ref. (69, 70)), we limited our search to multicellular organisms. We removed outgroups and duplicated species by hand and ensured that the trees were ultrametric and had branch lengths in units of millions of years. Tree manipulations and adjustments were done with the R packages ape (55), phangorn (56), castor (54), and rncl (53).

#### Estimating diversification rates from phylogenetic data

We calculated diversification rates (speciation and extinction) for each tree using BAMM v2.5.0 (21). The BAMM model is a variety of a birth-death process where the rates vary both through time and across the tree. BAMM uses reversible-jump MCMC (71) to locate breakpoints in the birth-death process on the phylogeny. We used BAMMtools (57) to configure the analysis; we used the default priors and incorporated a sampling fraction for each tree (using the true diversity estimated from the literature, as above) to correct the diversification rate estimates (59). For each tree, we ran 4 independent chains of 2 million generations each. We assessed mixing using the *effectiveSize* (ESS) function in the coda package (58). In cases, where the ESS was less than 100, we continued running the analysis until convergence was reached. For each of the 104 analyses, we computed the following summary statistics from the posterior distribution: i) the mean speciation rate across all branches (Fig. 1); ii) mean extinction rate across all branches; iii) speciation and extinction rates variance across all branches; and iv) the frequency of well-supported shifts (as identified by the protocol in BAMMtools (57)). We estimated the number of detectable shifts (>1) given clade age, and the average waiting times between evolutionary regime changes (*SI Appendix*, Fig. S1).

The standard BAMM model (21) fit to a tree with *n* tips includes *m* evolutionary regimes, where *m* can vary between 1 (single rate) and 2*n*-3 (each branch has its own rate). Each of the evolutionary regimes contains its own speciation and extinction rate, as well as a rate *k* the describes how the diversification process changes through time. The *k* parameter was included in the BAMM model owing to apparent widespread inadequacy of constant rate birth death models (72). A negative *k* reflects a slowdown in diversification (perhaps owing to diversity dependent diversification (27, 73), or to other processes and artifacts (74, 75) whereas a positive value of *k* indicates a speedup. As such, the estimate of the *k* values within a tree can indicate whether the time-dependency of rates across trees reflect true diversification trends. While a *k* parameter is estimated for each evolutionary regime, it is difficult to meaningfully summarize this across the tree (D. Rabosky, pers. comm.). We therefore took an alternative approach: we conducted another BAMM analysis on each tree (following the protocol described above), but this time constrained the model to include only a single evolutionary regime and calculated the mean *k* value across each posterior (Fig. 4). In addition to *k* we also computed the γ-statistic (76), another metric of changing diversification rates through time, both in its standard form and correcting for tree size (77).

BAMM has been criticized as an inappropriate model for diversification analyses (78) (though these critiques are themselves potentially problematic; see refs. (79, 80)). However, as we are not primarily interested in detecting the number and location of shifts (the focus of the critiques) we have little reason to believe that our estimates of mean rates are in any way biased. In any case, as an additional check, we computed diversification rates for each phylogeny using two alternative methods: i) the maximum likelihood estimator (MLE) for a constant rate model including incomplete sampling (59, 81), as implemented in diversitree (62); and ii) the mean estimate of the DR statistic (32) across all species in the tree.

We fit linear models on a log-log scale between mean rates (speciation and extinction estimated from BAMM) and crown ages extracted from the from tree heights (results in main text). Time-dependency explains 34% of variation in diversification rates across the tree of life (*R*^*2*^ value from fitting a linear model between the mean diversification rate [speciation - extinction] for each group and its crown age). We fit the same linear regression models using the MLE rates (*SI Appendix*, Fig. S2) and for the DR statistic (*SI Appendix*, Fig. S3) with qualitatively consistent results.

#### Curation of fossil time series

For the mammalian data, we used the data from Sugawara and Quental (in review). The authors of this study downloaded all occurrences associated with each of the 21 Placentalia orders (i.e., Afrosoricida, Artiodactyla, Carnivora, Cetacea, Chiroptera, Cingulata, Dermoptera, Erinaceomorpha, Hyracoidea, Lagomorpha, Macroscelidea, Perissodactyla, Pholidota, Pilosa, Primates, Proboscidea, Rodentia, Scandentia, Sirenia, Soricomorpha, Tubulidentata) in the Paleobiology Database (155; https://paleobiodb.org/) and in the New and Old Worlds - Database of fossil mammals (NOW; http://pantodon.science.helsinki.fi/now/). After checking for typographical errors and synonyms, Sugawara and Quental removed occurrences that were duplicated in the PDBD and NOW. The mammalian record is very well curated and the taxonomic information is more complete than the plants. The data includes the number of times each genus (the basic unit of paleobiological analyses) was recorded as well as the geological stage of each record. We conducted all analyses at the order level and restricted our dataset to genera with at least two records and orders that had at least 10 genera. In total, this resulted in a dataset including 16 orders of placental mammals (Artiodactyla, Carnivora, Cetacea, Chiroptera, Cingulata, Erinaceomorpha, Hyracoidea, Lagomorpha, Macroscelidea, Perissodactyla, Pilosa, Primates, Proboscidea, Rodentia, Sirenia, Soricomorpha), which included a total of 107,376 occurrences (median number of occurrences per order: 2,260.5) distributed in 3,180 mammalian genera (median number of genera per order: 73.5).

The paleobotanical datasets in the 155 are not as well curated as the mammalian data so an additional set of data cleaning steps were required. To do this we modified the protocol of ref. (82). First, we downloaded (April, 2018) all fossil occurrences for the following groups, individually: Arberiopsida, Bennettitopsida, Cladoxylopsida, Cycadopsida, Dicotyledonae, Equisetopsida, Filicopsida, Ginkgoopsida, Lycopsida, Magnoliopsida, Peltaspermopsida, Pinopsida, Polypodiopsida, Progymnospermopsida, Psilophytopsida, Pteridopsida, Voltziopsida, Zosterophyllopsida, Alismales, Apiales, Arales, Arecales, Asterales, Bennettitales, Buxales, Callistophytales, Caryophyllales, Caytoniales, Celastrales, Ceratophyllales, Coniferales, Cooksoniales, Cordaitales, Cordaitanthales, Cornales, Corystospermales, Cyatheales, Cycadales, Cyperales, Dilleniales, Dioscoreales, Dipsacales, Equisetales, Ericales, Euphorbiales, Fabales, Fagales, Filicales, Gentianales, Geraniales, Gleicheniales, Glossopteridales, Gondwanostachyales, Isoetales, Juglandales, Lamiales, Laurales, Magnoliales, Malpighiales, Malvales, Marsileales, Myrtales, Myrtiflorae, Nelumbonales, Nymphaeales, Osmundales, Peltaspermales, Pinales, Polygonales, Polypodiales, Proteales, Pseudosporochnales, Psilophytales, Ranunculales, Rhamnales, Rosales, Rubiales, Salviniales, Sapindales, Sapotales, Saxifragales, Schizaeales, Scrophulariales, Solanales, Sphenophyllales, Theales, Triuridales, Typhales, Umkomasiales, Urticales, Violales, Voltziales and Zingiberales.

We then removed duplicate entries filtering by the “occurrence_no” and checked the generic names against the Index Nominum Genericorum (ING;(83)). We then assigned each genus to a family and order using the following protocol. First, we checked if each genus was listed by the Pteridophyte Phylogeny Group (84), which contains all Pteridophyte extant genera and their respective higher-level classification. If it was not included in the list, then we checked if at least two databases (155, ING, TPL) included a family name for a genus and the name was the same across all the databases, we assigned that genus to that family. Then we cross-referenced the family name with The Plant List (TPL) v1.1 http://www.theplantlist.org/. If this was not the case, we searched three databases that aggregated taxonomic information: Encyclopedia of Life (EOL; http://eol.org/), Integrated Taxonomic Information System (ITIS; https://www.itis.gov/) and Global Biodiversity Information Facility (GBIF; https://www.gbif.org/). In cases of conflict between databases, we chose the most recent evidence-based taxonomy. After we assigned the Family name, we checked both PPG I 2016 and the Angiosperm Phylogeny Group (85) to assign the Family to its corresponding Order. For the genera that we could not assign a Family name or the Family name was not found in the Phylogeny Groups (84, 85), we used the most recent evidence-based information about Order in the queried databases (i.e., 155, ING, TPL, EOL, ITIS, GBIF). The Family name was used to help us identify the corresponding order. As for mammals, all analyses were conducted at the ordinal level.

The final dataset contains 17,640 plant fossil occurrences (median number of occurrences per order: 570), distributed in 430 genera (median number of genera per order: 17) and 19 orders (Arecales, Cycadeoideales, Ericales, Fabales, Fagales, Lamiales, Laurales, Malpighiales, Malvales, Myrtales, Pinales, Polypodiales, Proteales, Ranunculales, Rosales, Sapindales, Saxifragales, Schizaeales, Selaginellales).

We also analysed the classical Sepkoski’s compendium of marine animals (86) (downloaded from http://strata.geology.wisc.edu/jack/). Which includes information about the first and last occurrences for 9 Phyla (i.e., Actinopoda, Annelida, Arthropoda, Brachiopoda, Bryozoa, Chaetognatha, Chordata, Ciliophora, Hemichordata, Hyolitha, Trilobozoa). After removing the orders with less than 10 genera, we are left with 2,851 genera (median number of genera per order: 32) and 51 orders (i.e., Nassellaria, Spumellaria, Agnostida, Asaphida, Bradioriida, Corynexochida, Decapoda, Eurypterida, Lichida, Metacopida, Odontopleurida, Palaeocopida, Phacopida, Podocopida, Proetida, Ptychopariida, Redlichiida, Acrotretida, Atrypida, Lingulida, Orthida, Pentamerida, Rhynchonellida, Spiriferida, Strophomenida, Terebratulida, Thecideida, Cheilostomata, Cryptostomata, Cyclostomata, Cystoporata, Fenestrata, Trepostomata, Alepisauriformes, Antiarcha, Arthrodira, Beryciformes, Chelonia, Coelocanthiformes, Conodontophorida, Crossognathiformes, Elopiformes, Lamniformes, Pachycormiformes, Palaeonisciformes, Petalichthida, Pteraspidomorphes, Rajiformes, Semionotiformes, Hyolithida, Orthothecida).

#### Estimating diversification rates from fossil time series

To be consistent with the paleobiological literature, we use the term origination rather than speciation as we are working with higher level taxonomic groups. Also, as pointed out by Ezard et al. (87), the dynamics of morphological divergence recorded by the fossil record may differ from those of lineage divergence. Many of the estimators of origination and extinction rates that have been proposed for the analysis of fossil time series are inherently dependent on interval length (88). This time dependency was the motivation for the development of the now ubiquitously used Per Capita (PC) rate estimators (25) as they, like the rate estimators used with phylogenetic data, explicitly control for time (88). The PC rate estimators use only the first and last occurrence of a lineage and bin all observations by geological interval. As with the other analyses in this paper, we summarized the rates by computing mean origination and extinction rates across all intervals over which a group occurred. Unlike the analyses of the molecular phylogenies, crown age was not synonymous with duration as some groups had gone extinct entirely; we therefore measured duration as the time between the very first and very last occurrence of each group.

As with phylogenies we wanted to explore if our results arise from an accelerating secular trend within clades towards the present. We estimated the correlation between the rate estimates and the time-bin ordination (i.e., we wanted to test whether bins later in the series tended to have higher or lower rates than bins earlier in the series). We do not observe a strong or consistent trend of decreasing (or increasing) rates through time (origination: Spearman’s ρ = 0.138; extinction: Spearman’s ρ = −0.183). In concordance with the results from the phylogenetic analyses (above; Extended Data Fig. 3), fossils do not show evidence of evolutionary speed-ups towards present nor systematic slowdowns. Conversely with phylogenetic results, fossil rates show a qualitatively similar time-dependency trend among calculated (mean or median) estimators (*SI Appendix*, Fig. S4)

For rates estimated we fit a linear regression model between the natural logarithm of the rates and the natural logarithm of the duration of the entire group.

#### Evaluating purely statistical explanations for our results

We considered a number of artifactual statistical explanations that might explain why macroevolutionary rates scale negatively with time. On their own, it appears that none of these factors are sufficient to explain our results, though we acknowledge that it is difficult to completely rule out that some complex interaction of statistical artifacts could generate the pattern; however, resolving this is beyond the scope of the present paper.

The first, and simplest statistical artifact is that the negative time-dependency simply reflects the fact that in order for a young group to be observed -- and to be large enough to be considered for inclusion in an analysis such as ours -- it must have had high rates of diversification early in its history. As clades get larger (and older) the rates tend to regress towards the mean, such that young groups would tend to have inflated estimates. This sampling artifact is known as the “push of the past” in the macroevolutionary literature and has been documented in both the paleobiological (89) and phylogenetic data (90, 91). (As an aside, we note that this is related to, but distinct from, the “pull of the present/recent”, the artifact resulting from the fact that extant taxa have not yet been pruned by extinction; in our data, extinction rates also scale negatively with time, which is inconsistent with the “pull of the present” artifact being an important part of the explanation.) As we show below, even when trees are simulated under a birth-death (speciation and extinction) or pure-birth (speciation only) process conditioned on surviving and containing a sufficient number of lineages, we observe a pattern where rates are inflated for lineages that originated close to the present (as in our empirical data). The key question is whether this effect is strong enough to explain our findings; using simulations designed following some recently developed theory (89), we are able to answer that it is not likely to be the case.

We estimated the mean rates of speciation and extinction from the oldest groups (more than 150 million years), where the curve between rates and age levels off (on a natural scale; Figure 2); this gave us parameters of λ = 0.11 and µ = 0.08. We then used TreeSim (61) to generate 104 phylogenies with the same ages as our empirical trees. We conditioned the simulations to produce trees with at least 7 taxa (the same as in our empirical trees) such that for younger clades, the only trees that would meet this condition were those that had stochastically high number of speciation events early in their history. We repeated this procedure 1000 times and estimated the slope of the log(speciation rate) ∼ log(crown age) regression resulting from each one. We then compared this to our empirically estimated slope. We found that the slopes generated by conditioning alone were indeed negative but not nearly as negative as our empirically estimated slope (*SI Appendix*, Fig. S6), suggesting that our empirical results do not simply reflect inflated estimates for young groups. As a further check, we repeated this procedure by setting µ = 0.5λ since extinction rates are notoriously difficult to estimate (92). This scenario was qualitatively similar to those from the empirically estimated extinction rate.

Second, the groups we have included in this meta-analysis are not random subsets of the Tree of Life; the very fact that they are generally named groups means that taxonomists have identified these as special or distinct in some way (93). Rabosky and colleagues (27, 28, 94) have suggested that some as-yet-unidentified artifact of taxonomic practice may break the correlation between age and species richness -- and consequently, generate a negative time-dependency of macroevolutionary rates. Even without a clear mechanism, we can test this proposition. To do this, we used recently published megaphylogenies of birds (32), ferns (40), and Angiosperms (41). We first computed the MLE of diversification rates for named clades (families, families, and orders, respectively) consisting of at least 10 taxa using the package diversitree (62) and fit a linear regression model log(speciation rate) ∼ log(crown age) through all of the subclades. We then randomly selected an equivalent number of nodes from each tree with (approximately) the same age distribution as the empirical named groups. We computed the MLE diversification rates for each of these subtrees and computed the same regression model as with the named nodes. We repeated this process 1000 times. Finally, as with the ascertainment bias simulations, we compared the regression slope from the named nodes to the distribution of slopes from random subtrees and in all cases, found no difference (Fig. 4). In other words, we see the same pattern of rate-scaling whether we use named clades or random nodes within the Tree of Life. This result strongly suggests that our negative slope results are not a sampling artifact due to taxonomic delimitation; there is nothing particularly special, at least in terms of diversification dynamics, between the groups descending from named nodes (e.g., those representing Linnean ranks) and unnamed nodes. (It is not possible to subset the fossil time series at unnamed clades so we assume that our results from the phylogenetic data can be applied to the fossil data as well.)

While we tried to be as systematic as possible in collecting trees from the literature so as not to bias our results, it is nonetheless possible that scientists themselves are biased in terms of what groups they build trees for; for example, they may tend to study groups that are more diverse than average. If this selection bias is more pronounced for younger groups, then it is possible that our negative time-dependency might reflect this. However, we cannot imagine any way to evaluate this with our data.

Third, many of the trees in our collection were from sparsely sampled groups (and the older groups tended to be more sparsely sampled than younger ones; Spearman’s rank correlation between age and sampling fraction: −0.25). While previously simulation studies (e.g., ref. (59)) have demonstrated that birth-death estimators are consistent when the correct sampling fraction has been provided, we performed a brief simulation of our own to ensure our results were not susceptible to our particular pattern of sampling. We used the empirically estimated ages and sampling fractions from our phylogenetic analyses and used the TreeSim R package (61) to simulate a tree corresponding to each empirical dataset (as above, using the mean rates estimated from empirical phylogenies older than 150 My; λ = 0.11, µ = 0.08). We re-estimated the MLE parameters using diversitree (62) and then fit a linear model between the (log) estimates and (log) crown age as we did in the empirical data. We repeated this procedure 1000 times and compare the estimated slopes to our empirically estimated slopes. There is no indication that our pattern of sampling could generate the negative relationship between rates and age that we observe in our empirical data (*SI Appendix*, Fig. S7). However, we assume throughout that sampling is randomly distributed across the tree; of course in real life, researchers are likely to preferentially sample some taxa (e.g., to ensure that a phylogeny contains representatives from all major groups) and such sampling schemes can mislead birth-death estimators if not accounted for (95–97). While we acknowledge that failing to account for nonrandom sampling may bias our results (though in which way, it is not entirely clear), given the breadth of our study, there is no way we could feasibly control for the pattern of sampling in each individual dataset.

Fourth, it is a well known statistical fact that measurement error in the explanatory variable will tend to negatively bias the estimate of the slope. Two previous simulation studies have found that diversification rates are generally fairly robust to errors in divergence time estimation (98, 99) (but see (20)). Still, error in ages of young clades could lead to pathological overestimates of diversification rates, potentially driving a negative relationship between rates and time. To investigate this, we repeated the simulations described previously to test the “push of the past” artifact. This time, we added error to clade ages. For each age, we drew a percentage error from a uniform distribution, and then added or subtracted the appropriate amount from the branch length. We ran a set of simulations where maximum error varied from 10% to 90%. Although altering the error did result in negative scaling of diversification rate with clade age, in no case was the slope as extreme as in our empirical data (*SI Appendix*, Fig. S8). In other words, error in branch length estimation cannot explain our results.

And last, the estimators themselves may be slightly biased in a way that previous simulation studies have missed; a central claim of our paper is that the estimators are not expected to show any time-dependency. We think that this is very unlikely to be driving the pattern. General birth-death estimators have been extremely well studied in the context of macroevolution and have been found to be consistent (e.g., (100)). The phylogenetic method used in this study (BAMM (101)) can accommodate high levels of heterogeneity in the diversification process (both across time, and across lineages), making them more likely to be adequate than simple, constant rate models.

Nonetheless, our results are qualitatively the same whether we use the fits of complex models or much simpler metrics (such as the DR for phylogenies (32)).

### Code and data availability

All scripts for data curation and analysis are available on GitHub at https://github.com/mwpennell/macro-sadler. Since we used only previously published (and publically available) phylogenetic and paleobiological data for this analysis, we will not re-publish the curated datasets; however, the final, curated datasets are available upon request.

**Fig. S1.**
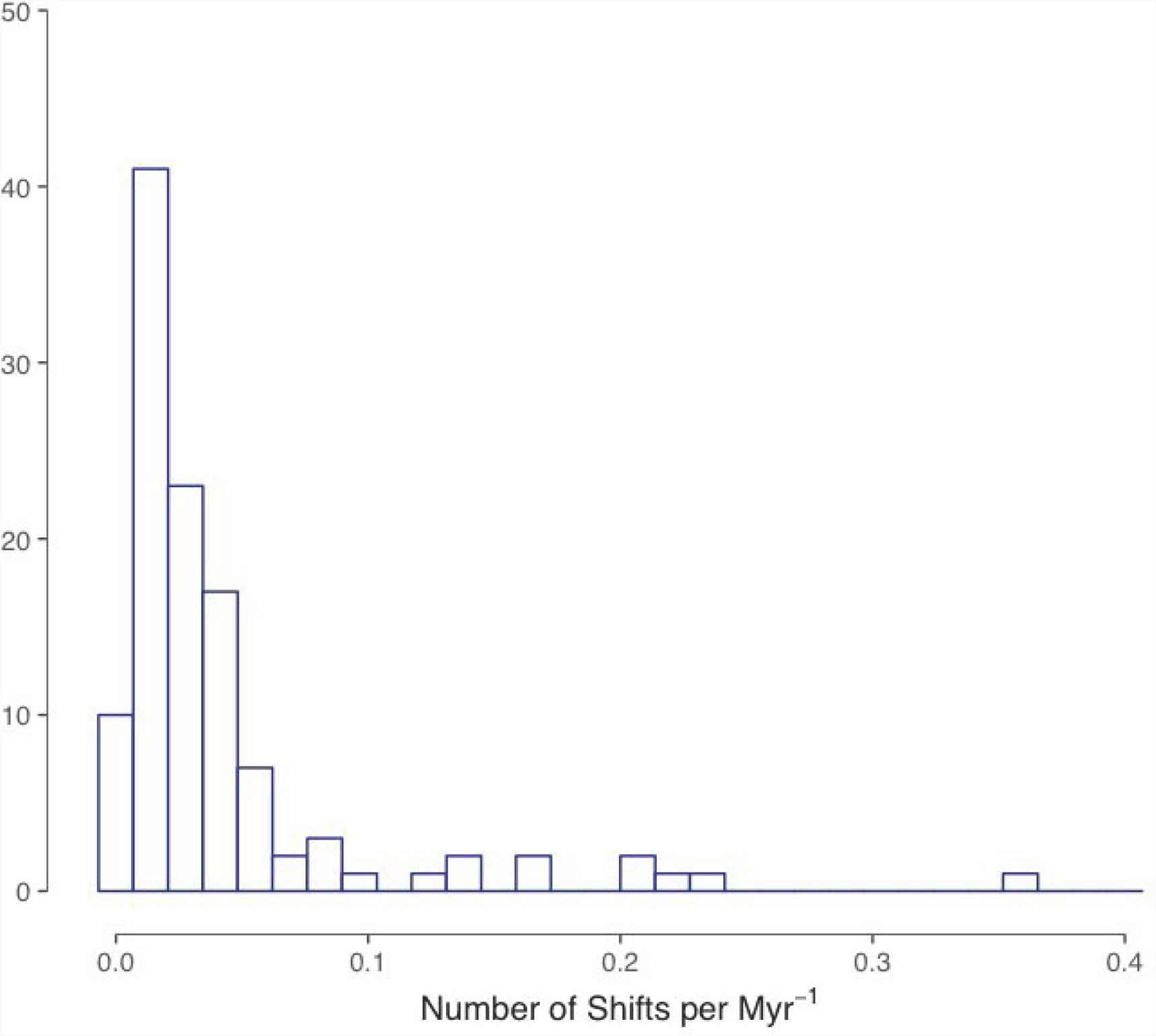
Frequency of number of highly supported shifts per million years of evolution. For each phylogeny we fit a reversible-jump BAMM model and used BAMMtools (57) to identify the “Best Number of Shifts” in each analysis. Since the number of highly supported shifts recovered depends highly on the number of taxa in the analysis (Spearman’s rank correlation rho: 0.51), we standardized the frequency by dividing my millions of years. On average, highly supported shifts occurred every 24.2 million years (variance: 0.03 My^2^). While it is almost certainly just a coincidence, it is noteworthy that this frequency is almost perfectly aligned with that of major shifts in phenotypic evolution (14, 102) and major extinction events observed in the fossil record (47, 103, 104). We will leave it to the reader to speculate about what this might mean.

**Fig. S2.**
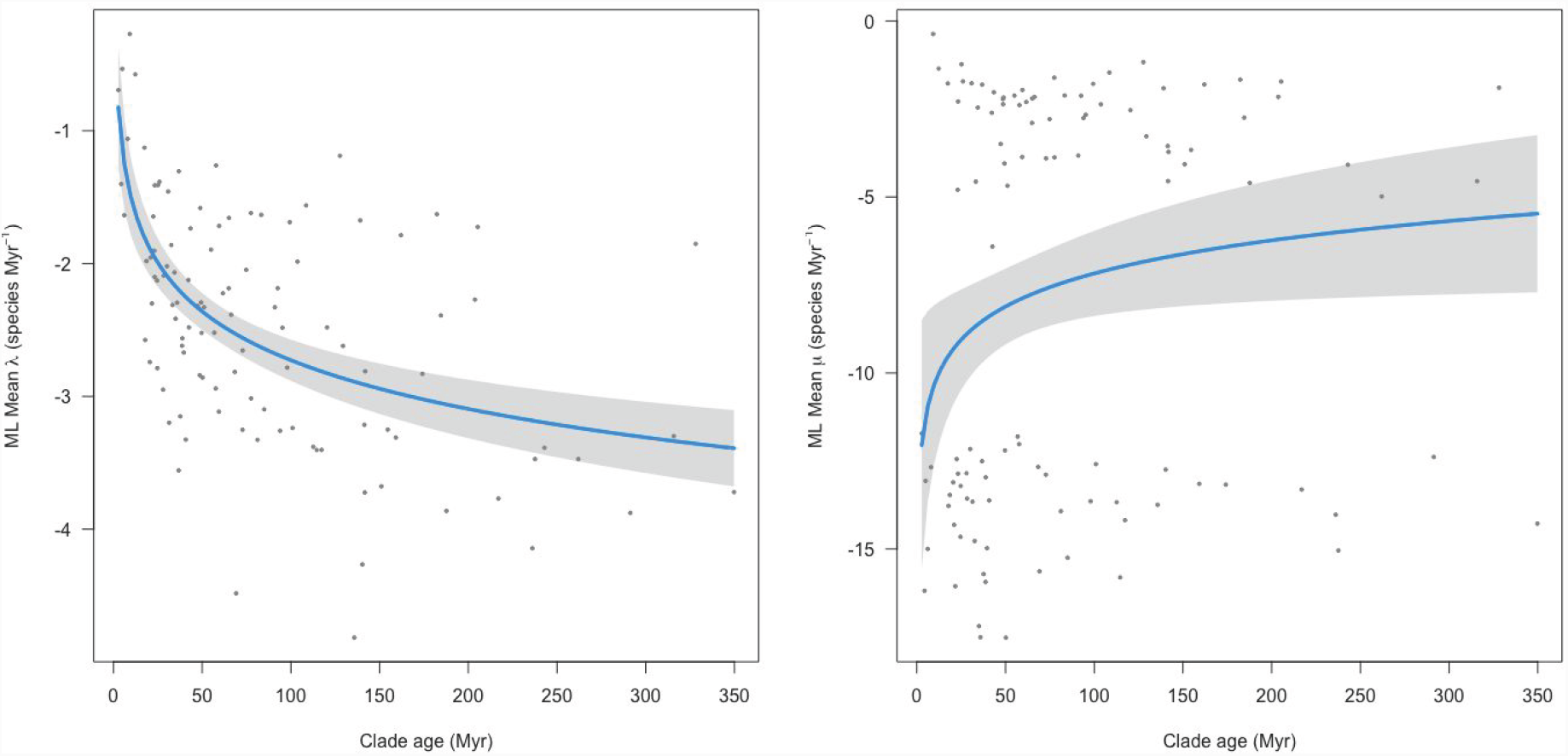
Maximum likelihood estimates of speciation (λ) and extinction rate (µ) vs clade age from our empirical data. The estimates have been corrected for incomplete sampling. As has been previously noted, MLEs for the extinction parameter are often unreliable, particularly when the sampling fraction is low (59, 92). As such, we do not put much stock in the lack of time-scaling in the extinction rates but include the plot for the sake of completeness.

**Fig. S3.**
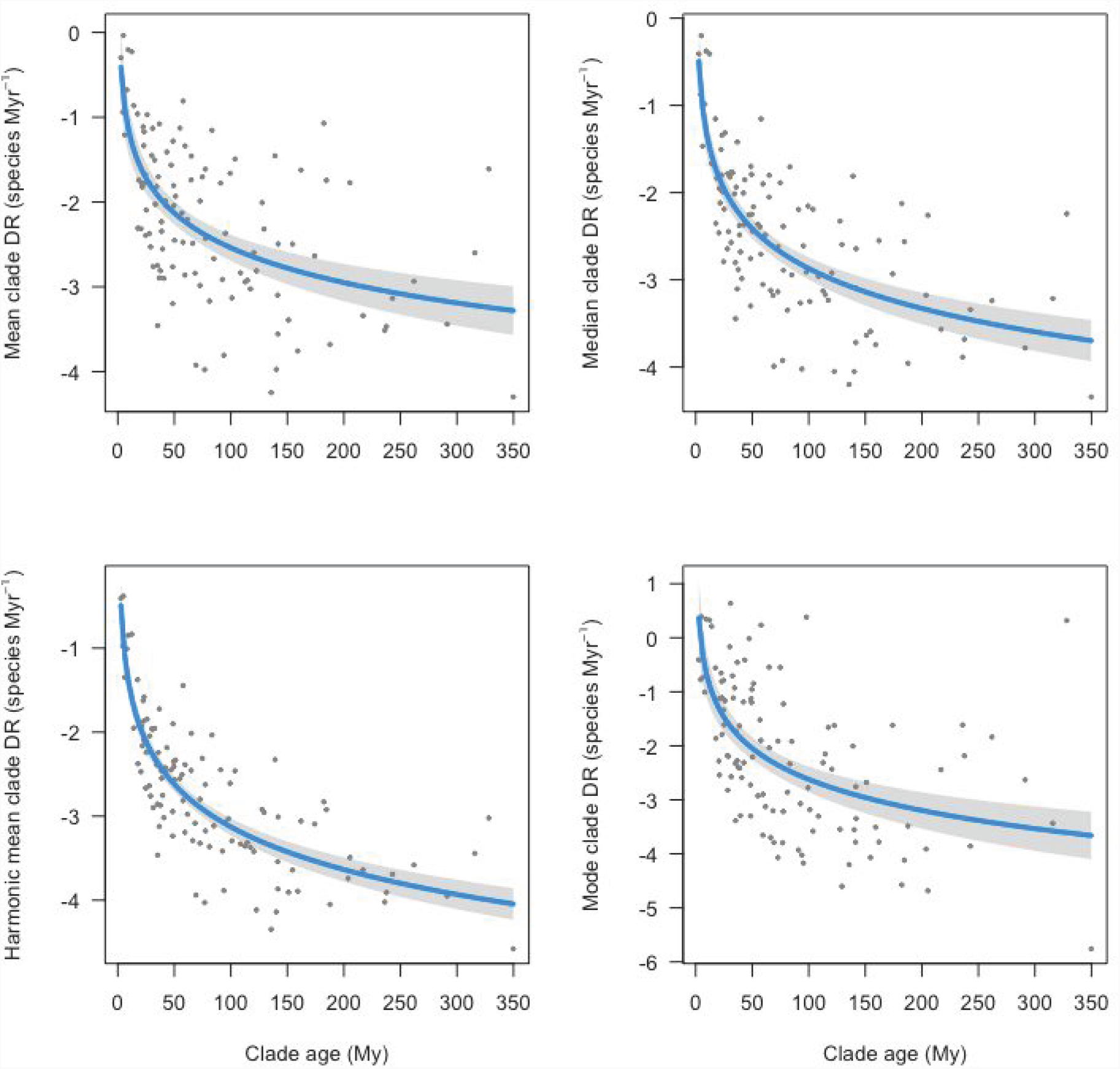
Tree-wide DR statistic vs clade age. As an alternative to the mean posterior rates estimated in the BAMM analyses, we also computed the DR statistic (32) for every tip and the mean, median, harmonic mean, and mode of the DR statistic for each tree. The average DR statistic, which is a non-parametric metric closely related to the speciation rate (32, 105), shows the same type of time-scaling as the rates estimated from BAMM; this demonstrates that the time-scaling is a general phenomenon and not specific to the BAMM model.

**Fig. S4.**
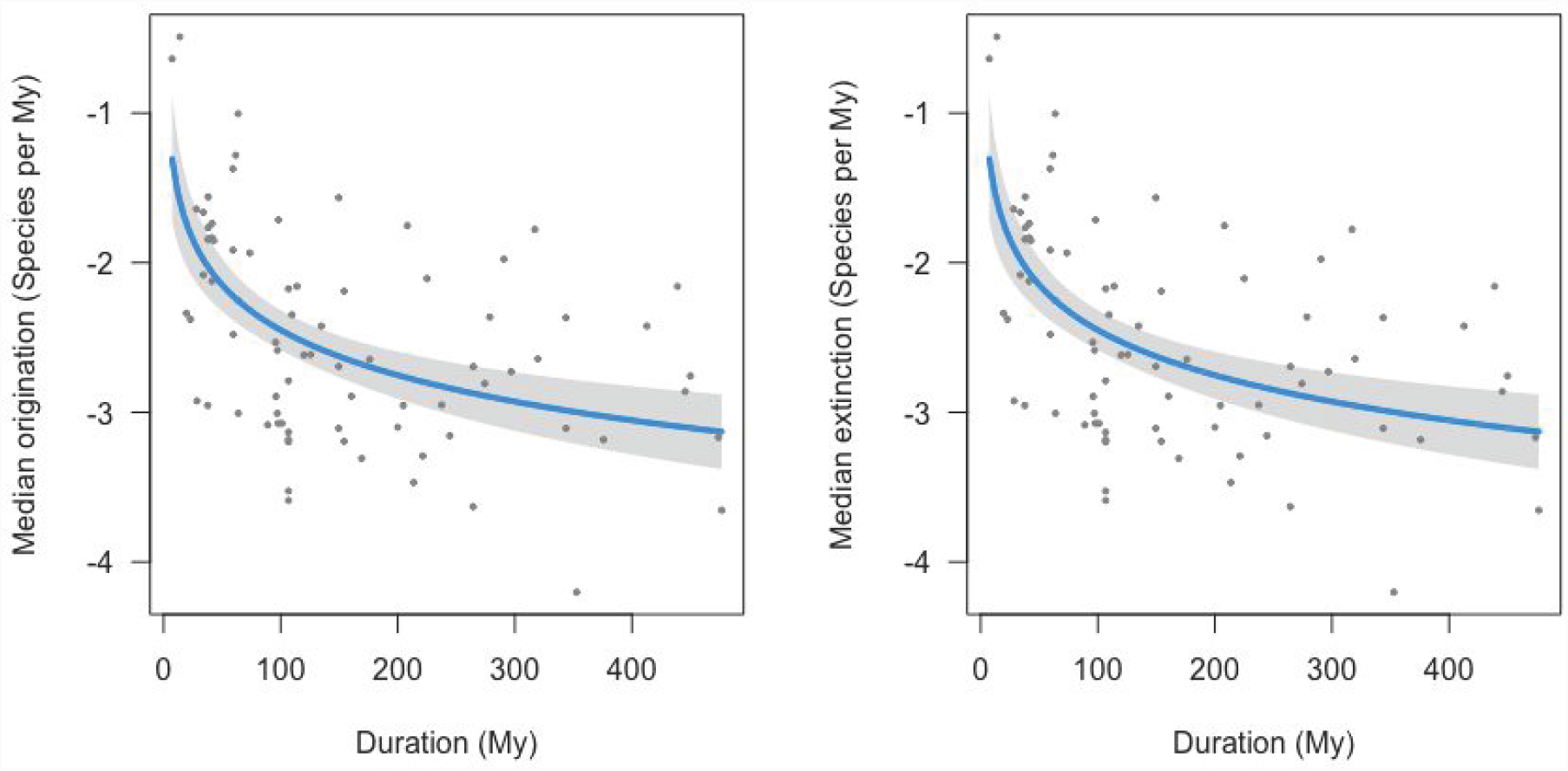
Median origination and extinction rates vs clade age (fossil data). Median rates were estimated for mammals, plants and Sepkoski’s marine animals estimated using the perCapita method. The similarity of the mean to the median demonstrates that the pattern of time scaling is not driven by boundary effects or outlying time intervals.

**Figure S5.**
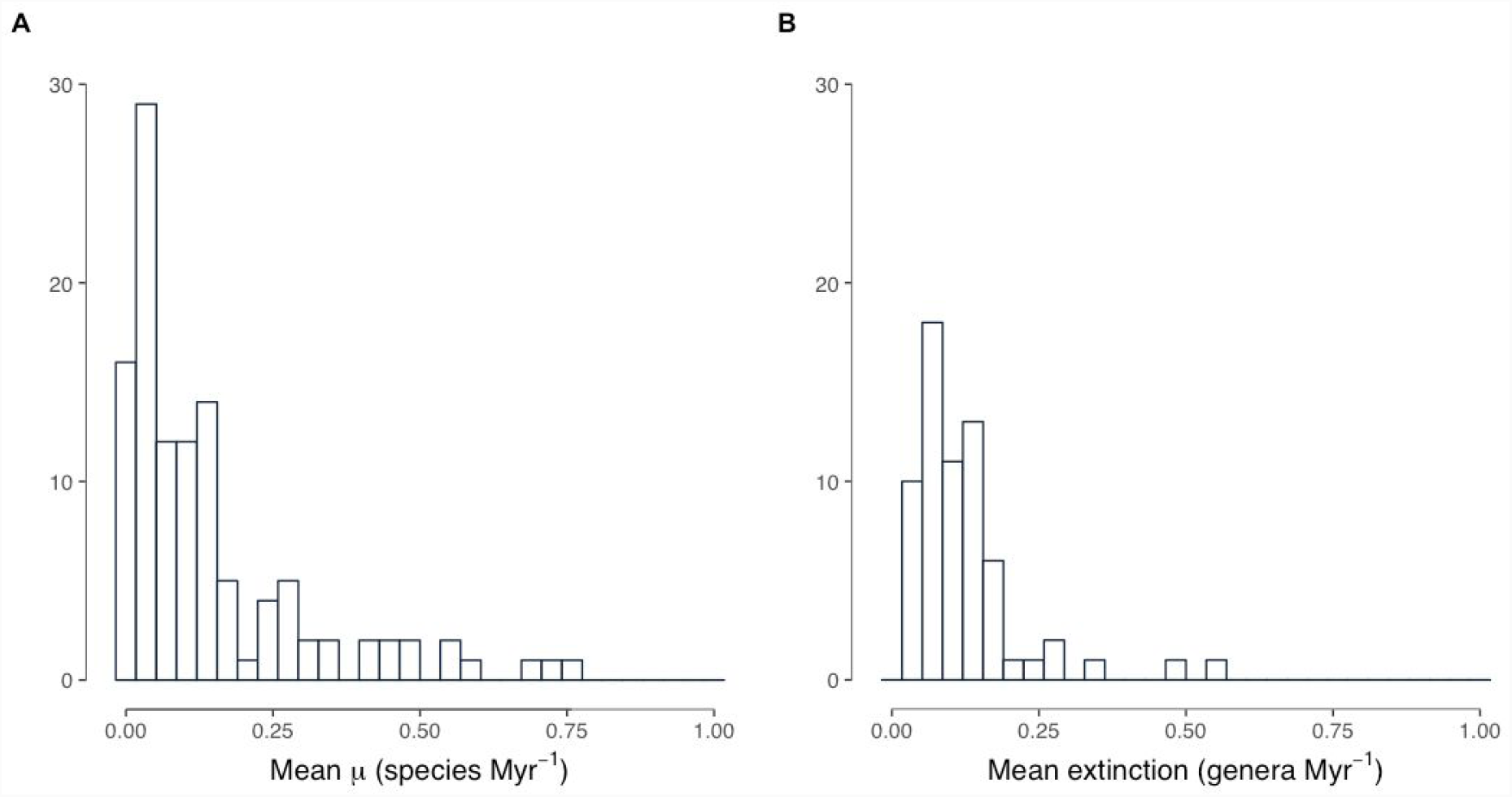
Histogram of mean extinction rates. A) The lowest extinction rate corresponds to the Elasmobranchii subclass (0.02), the highest is the bird family Psophiidae (0.76). B) The lowest extinction rate corresponds to the Mammal order Macroscelidea (0.02), the highest is Berciformes order (0.48).

**Fig. S6.**
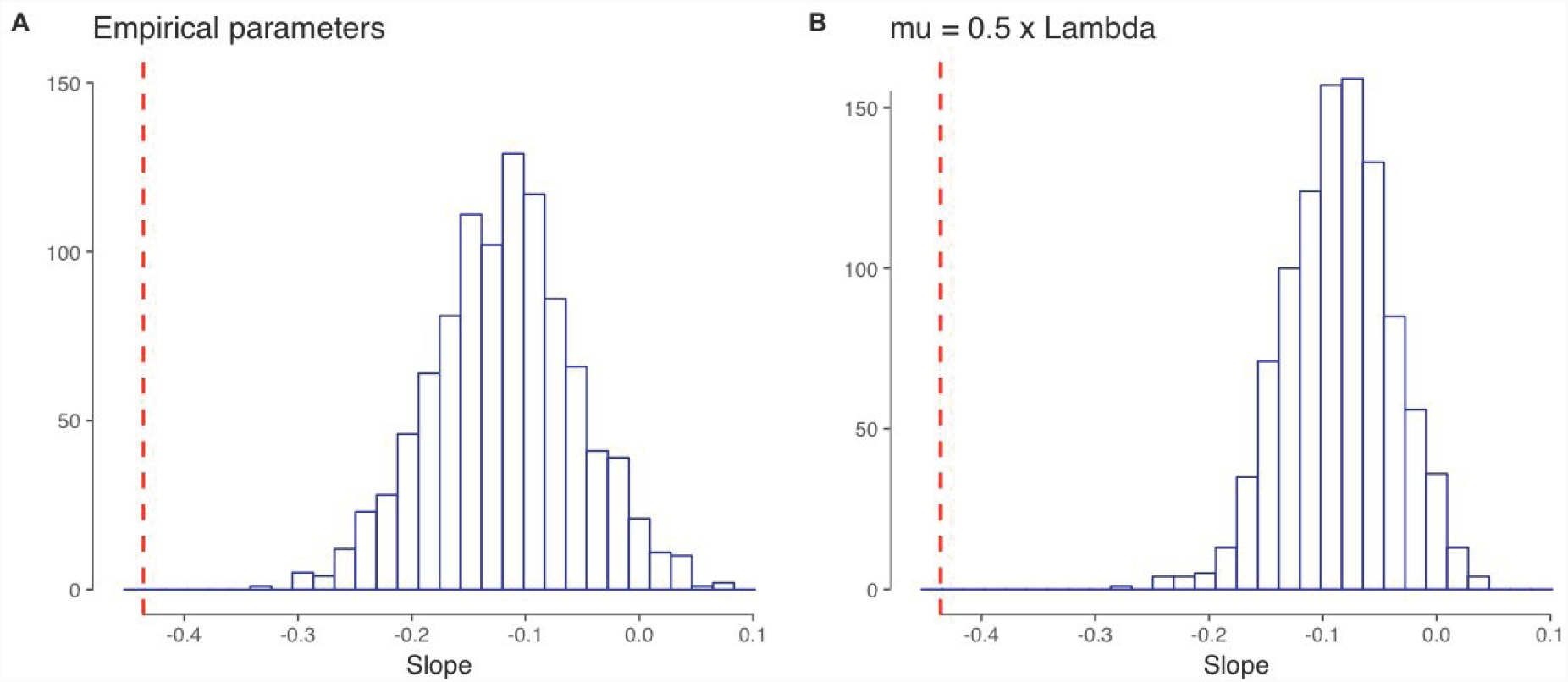
Push of the past effect. Empirical slope (dash) compared with slope’s distribution from simulated trees (parameters from trees older that 150 Myr.) with λ = 0.11 and differential extinction rates A) µ = 0.08 and B) µ = 0.5λ. Although the push of the past may generate negative slopes, our empirical slope are far less negative than those in our empirical and alternative scenario discarding it as our results main driver.

**Fig. S7.**
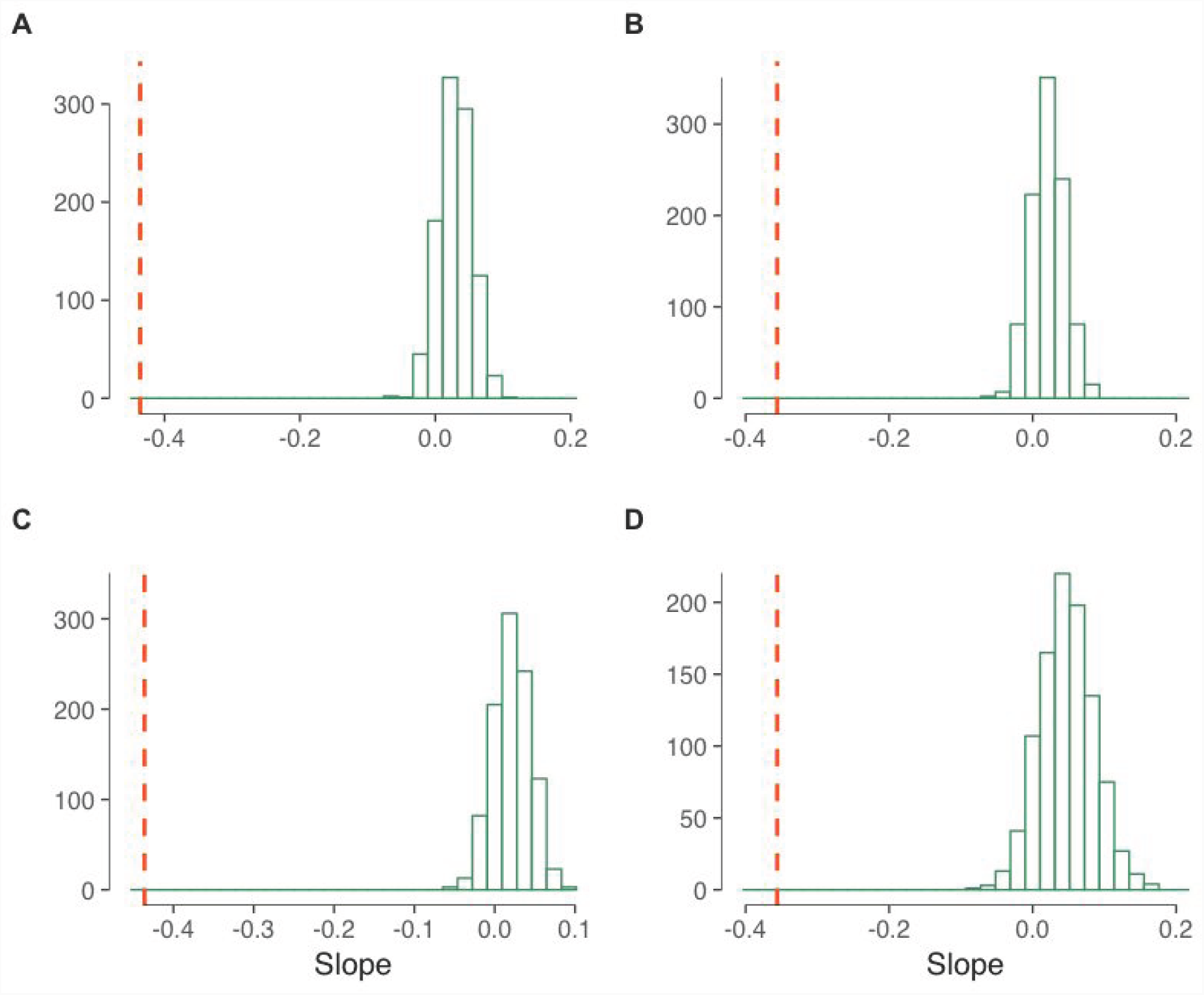
Sampling fraction effect. Empirical slope (dashed) compared with slope’s distribution from simulated trees. Speciation (A, C) and extinction rates (B, D). A) and B) Trees simulated with empirical sampling fractions; C and D) Trees simulated with empirical sampling fraction and age. We did not found any significative trend in any of our simulation scenarios nor speciation or extinction rates; if any it seems to be rather mainly positive. This suggest that in spite of sampling fraction effect in rates estimation it acts in a different direction than our negative dependent results.

**Fig. S8.**
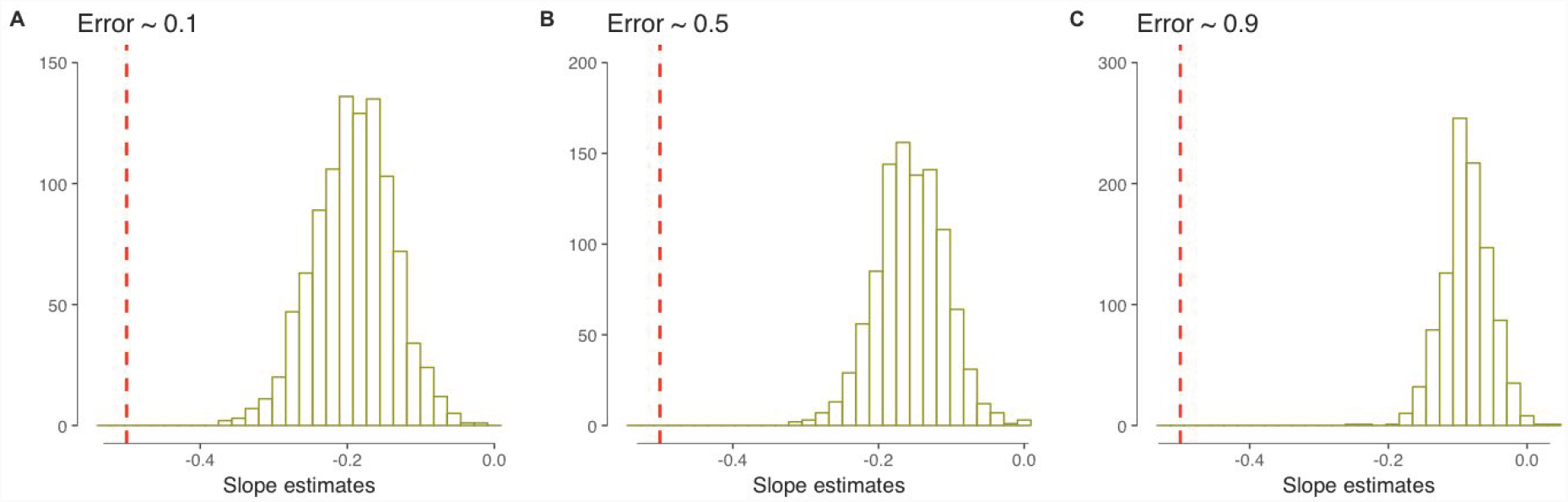
Branch length error effect. Empirical slope (dashed) compared with slope’s distribution from biased or error in tree’s branch lengths. Three different error scenarios A) 0.1, B) 0.5 and C) 0.9

**Table S1.**
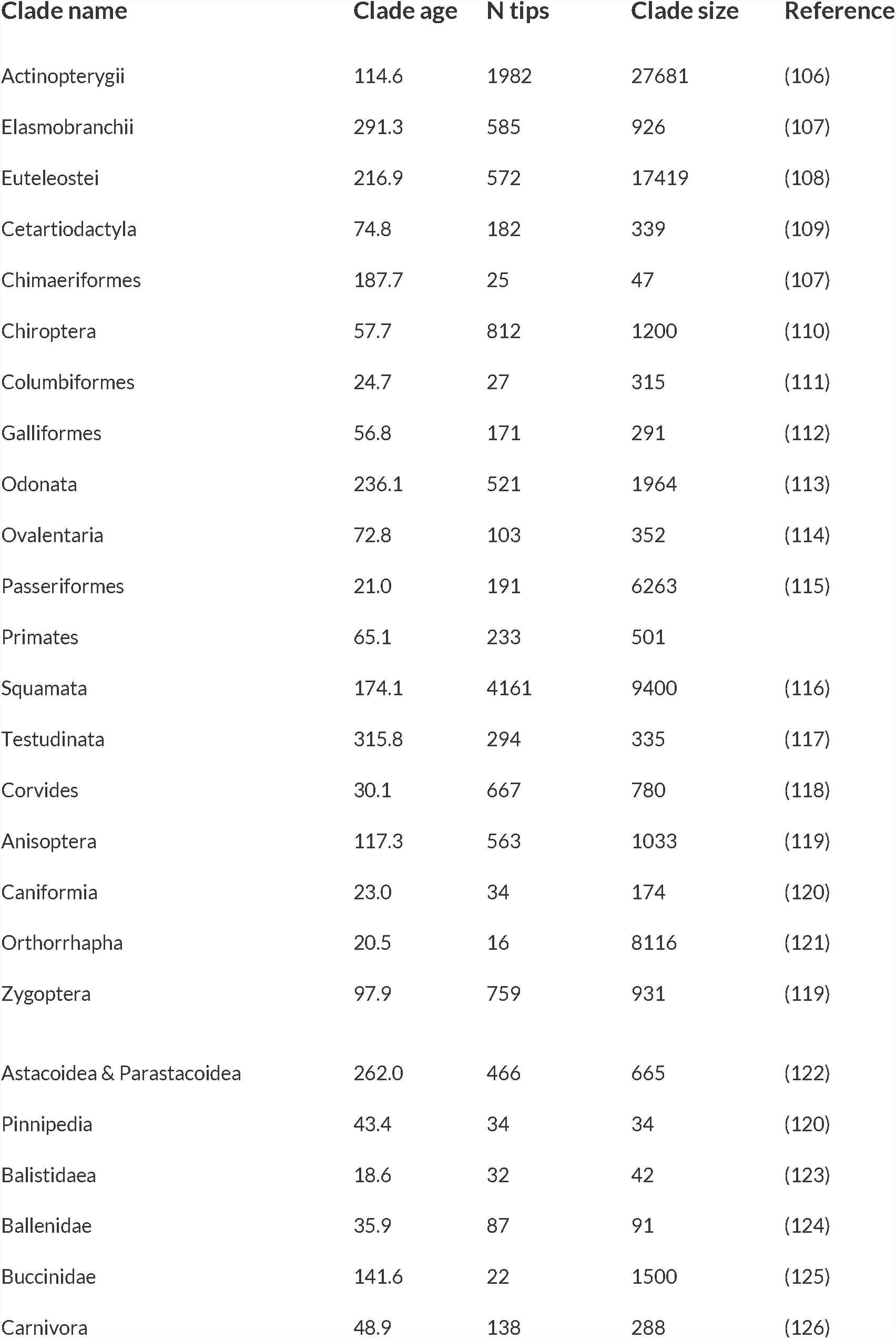
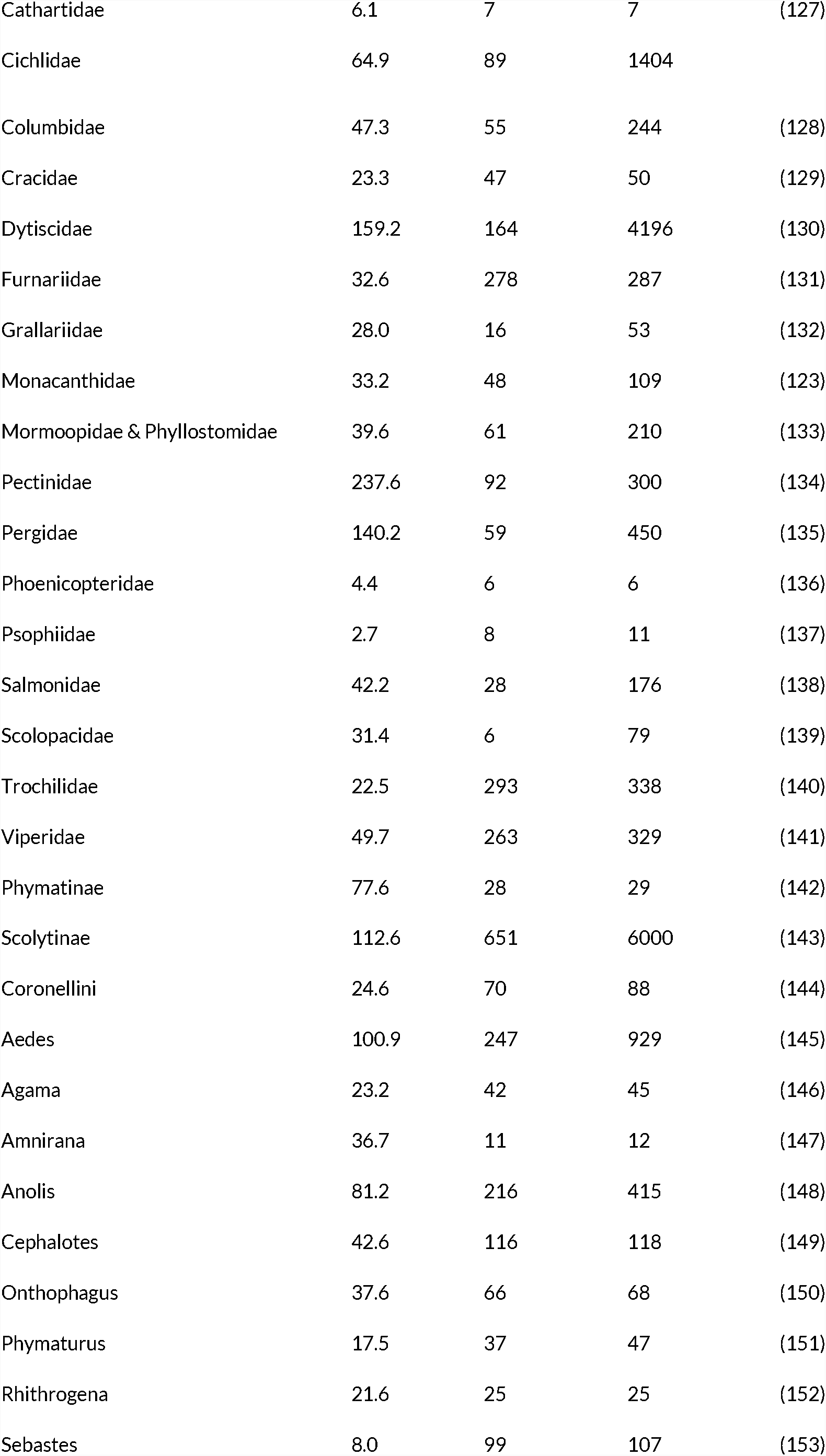
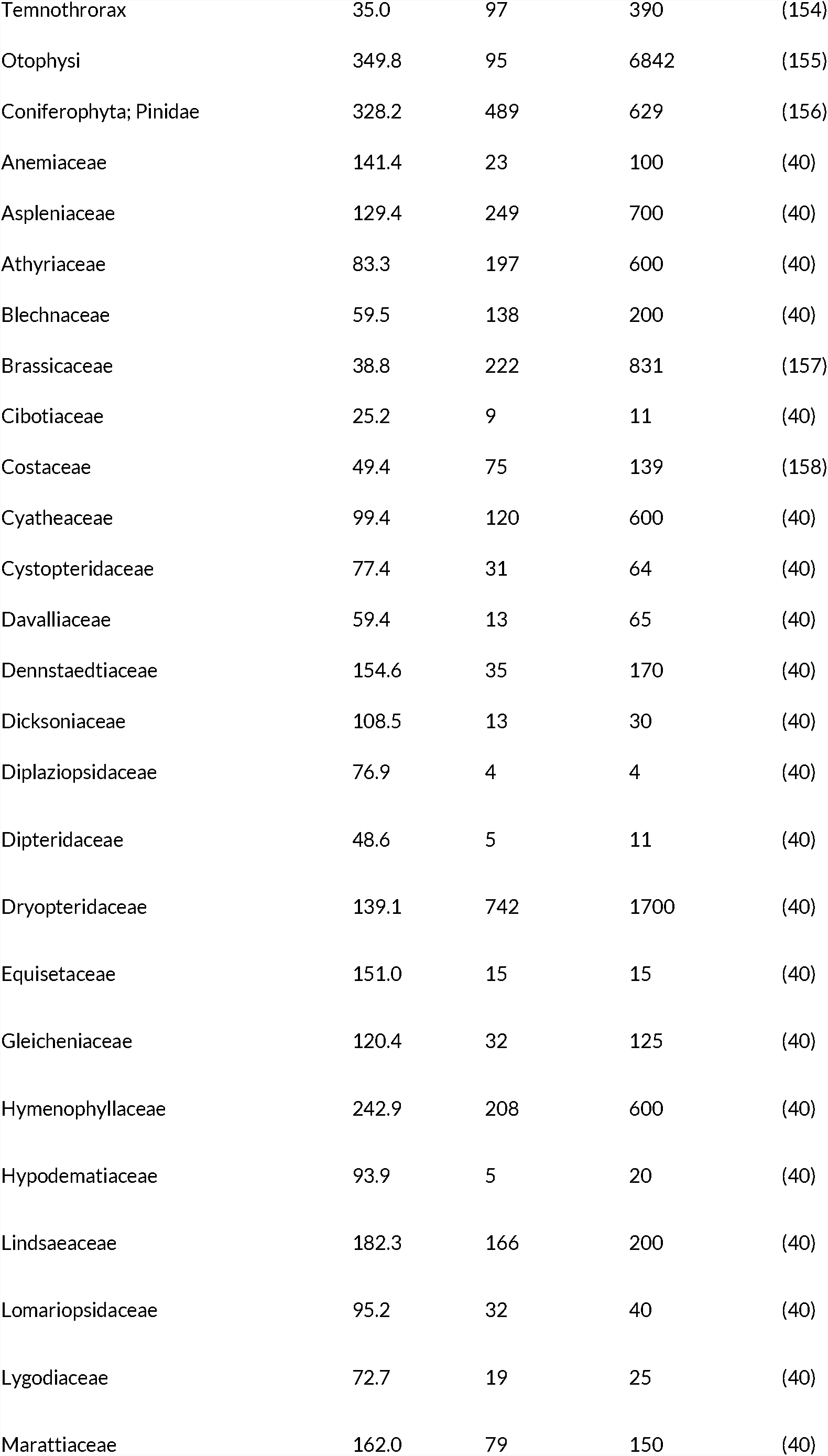
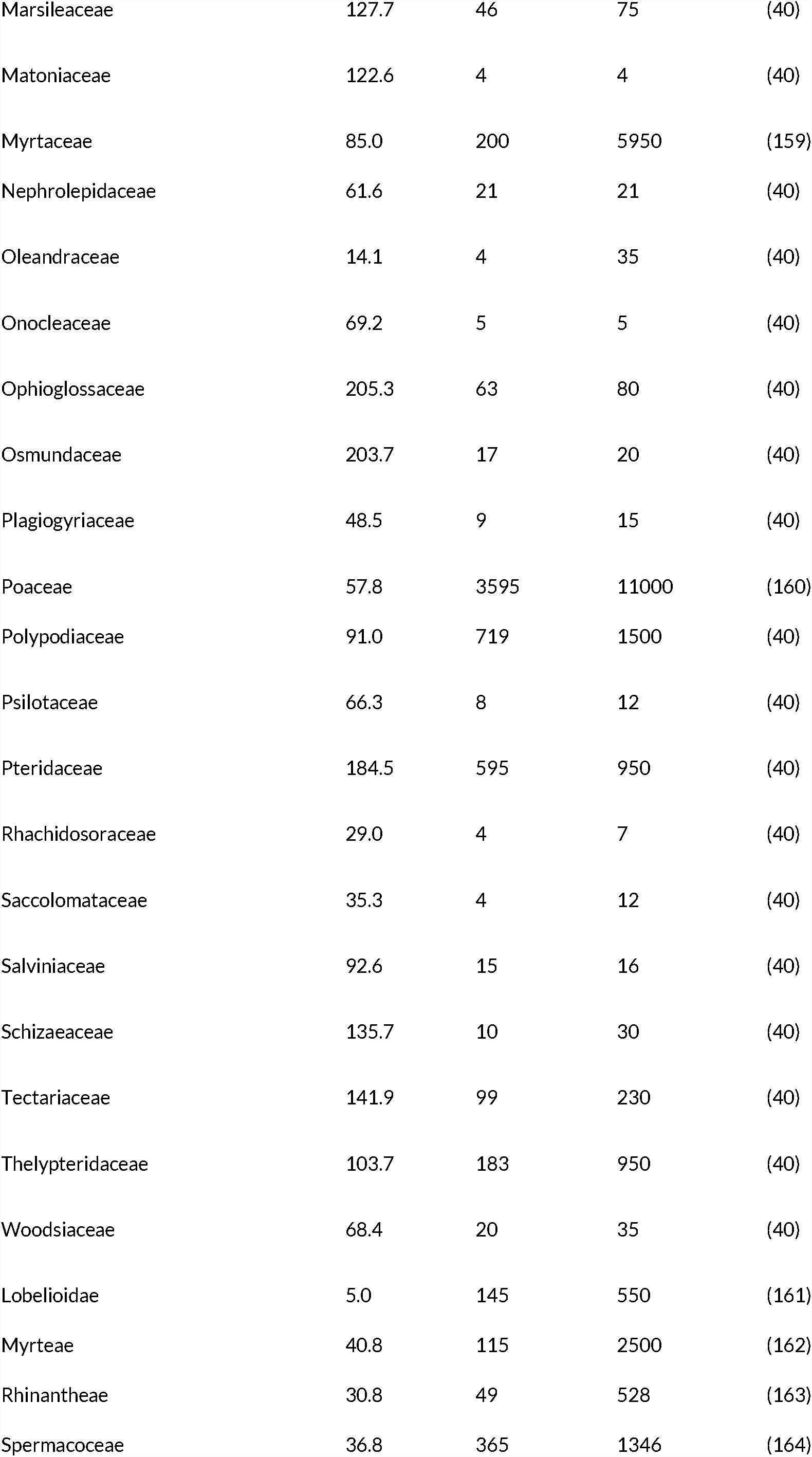
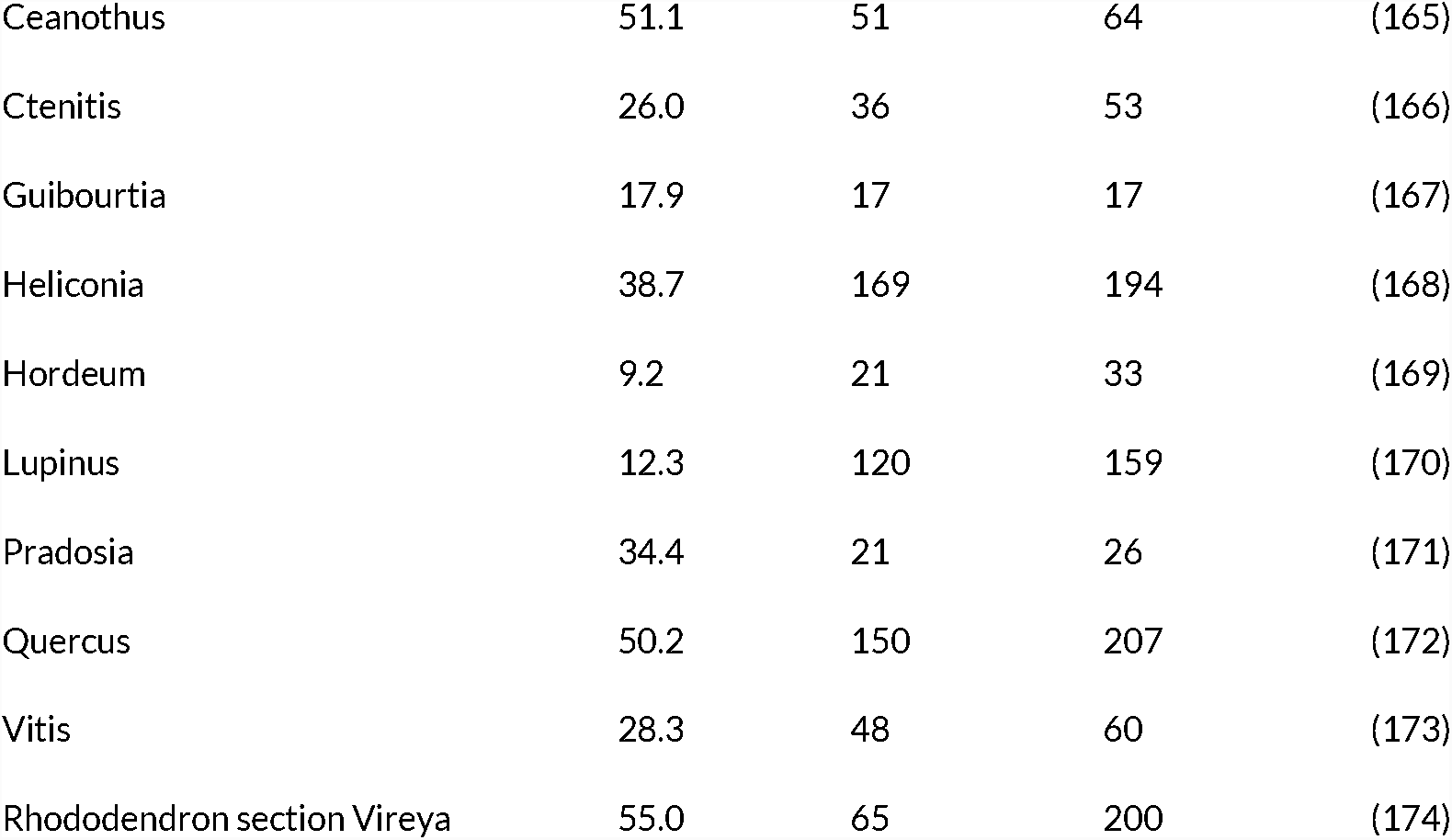
Summary of phylogenies used in this analysis.

**Table S2.** Summary of fossil dataset used in this analysis. “Duration” are shown in millions of years. Note: there is no information about the number of occurrences Sepkoski (2002) used to assign first and last appearances for each genus in the compendium.

| Clade name | Duration | Genera | Occurrences | Source |
| --- | --- | --- | --- | --- |
| Afrosoricida | 2.588 | 12 | 49 | (175, 176) |
| Artiodactyla | 38 | 552 | 11724 | (175, 176) |
| Carnivora | 41.3 | 456 | 7578 | (175, 176) |
| Cetacea | 33.9 | 351 | 2130 | (175, 176) |
| Chiroptera | 33.9 | 109 | 830 | (175, 176) |
| Cingulata | 7.246 | 50 | 351 | (175, 176) |
| Erinaceomorpha | 38 | 41 | 233 | (175, 176) |
| Hyracoidea | 23.03 | 24 | 156 | (175, 176) |
| Lagomorpha | 28.1 | 58 | 1703 | (175, 176) |
| Macroscelidea | 38 | 28 | 562 | (175, 176) |
| Perissodactyla | 59.2 | 225 | 6422 | (175, 176) |
| Pilosa | 13.82 | 37 | 323 | (175, 176) |
| Primates | 59.2 | 264 | 2726 | (175, 176) |
| Proboscidea | 41.3 | 50 | 2316 | (175, 176) |
| Rodentia | 41.3 | 807 | 14491 | (175, 176) |
| Sirenia | 38 | 41 | 391 | (175, 176) |
| Soricomorpha | 61.6 | 81 | 970 | (175, 176) |
| Acrotretida | 97.95 | 25 | NA | (177) |
| Agnostida | 127.25 | 35 | NA | (177) |
| Alepisauriformes | 28.5 | 13 | NA | (177) |
| Antiarcha | 46.75 | 10 | NA | (177) |
| Arthrodira | 42.25 | 28 | NA | (177) |
| Asaphida | 63.65 | 60 | NA | (177) |
| Atrypida | 73.6 | 53 | NA | (177) |
| Beryciformes | 19.5 | 22 | NA | (177) |
| Bradioriida | 42.85 | 25 | NA | (177) |
| Cheilostomata | 95.65 | 67 | NA | (177) |
| Chelonia | 149.7 | 15 | NA | (177) |
| Coelocanthiformes | 317.05 | 14 | NA | (177) |
| Conodontophorida | 268.4 | 16 | NA | (177) |
| Corynexochida | 149.6 | 84 | NA | (177) |
| Crossognathiformes | 59.45 | 17 | NA | (177) |
| Cryptostomata | 200 | 38 | NA | (177) |
| Cyclostomata | 445.45 | 78 | NA | (177) |
| Cystoporata | 264.4 | 40 | NA | (177) |
| Decapoda | 221.35 | 63 | NA | (177) |
| Elopiformes | 81.4 | 10 | NA | (177) |
| Eurypterida | 176 | 32 | NA | (177) |
| Fenestrata | 213.65 | 42 | NA | (177) |
| Hyalithida | 101.05 | 21 | NA | (177) |
| Lamniformes | 134.75 | 11 | NA | (177) |
| Lichida | 88.95 | 24 | NA | (177) |
| Lingulida | 438.85 | 18 | NA | (177) |
| Metacopida | 319.75 | 19 | NA | (177) |
| Nassellaria | 237.45 | 37 | NA | (177) |
| Odontopleurida | 113.95 | 11 | NA | (177) |
| Orthida | 149.6 | 88 | NA | (177) |
| Orthothecida | 134.75 | 11 | NA | (177) |
| Pachycormiformes | 154.2 | 28 | NA | (177) |
| Palaeocopida | 343.8 | 113 | NA | (177) |
| Palaeonisciformes | 343.75 | 51 | NA | (177) |
| Pentamerida | 125.6 | 59 | NA | (177) |
| Petalichthida | 25.05 | 10 | NA | (177) |
| Phacopida | 119.8 | 60 | NA | (177) |
| Podocopida | 476.6 | 130 | NA | (177) |
| Proetida | 225 | 71 | NA | (177) |
| Pteraspidomorphes | 97.6 | 27 | NA | (177) |
| Ptychopariida | 149.6 | 170 | NA | (177) |
| Rajiformes | 98.15 | 17 | NA | (177) |
| Redlichiida | 24 | 18 | NA | (177) |
| Rhynchonellida | 449.95 | 274 | NA | (177) |
| Semionotiformes | 154.2 | 12 | NA | (177) |
| Spiriferida | 274.2 | 213 | NA | (177) |
| Spumellaria | 473.95 | 52 | NA | (177) |
| Strophomenida | 278.6 | 227 | NA | (177) |
| Terebratulida | 412.7 | 229 | NA | (177) |
| Thecideida | 208.1 | 14 | NA | (177) |
| Trepostomata | 264.4 | 49 | NA | (177) |
| Cycadeoideales | 204.7 | 16 | 517 | (175) |
| Saxifragales | 169.231 | 11 | 290 | (175) |
| Cladoxylales | 37.7 | 12 | 73 | (175) |
| Cycadales | 290.661 | 11 | 220 | (175) |
| Fagales | 106.681 | 23 | 1489 | (175) |
| Ericales | 97.131 | 22 | 352 | (175) |
| Malpighiales | 106.681 | 24 | 456 | (175) |
| Rosales | 106.681 | 28 | 726 | (175) |
| Gentianales | 63.731 | 10 | 48 | (175) |
| Oxalidales | 63.731 | 11 | 111 | (175) |
| Malvales | 160.331 | 14 | 280 | (175) |
| Schizaeales | 375.762 | 16 | 552 | (175) |
| Polypodiales | 296.881 | 14 | 210 | (175) |
| Lycopodiales | 244.531 | 10 | 155 | (175) |
| Myrtales | 97.131 | 16 | 308 | (175) |
| Sapindales | 106.681 | 29 | 424 | (175) |
| Proteales | 106.681 | 34 | 592 | (175) |
| Laurales | 106.681 | 14 | 247 | (175) |
| Ranunculales | 106.681 | 12 | 205 | (175) |
| Fabales | 95.909 | 18 | 85 | (175) |
| Arecales | 109.562 | 13 | 295 | (175) |
| Pinales | 352.731 | 73 | 2892 | (175) |

**Table S3.** Time intervals of systematically reviewed Journals.

| **Journal** | **Last issue reviewed** | **First issue reviewed** |
| --- | --- | --- |
| Am. J. Bot. | Vol. 105 (1); 2018 | Vol. 103 (12); 2016 |
| BMC Evol. Biol. | Vol. 18:26; 2018 | Vol. 17:248; 2017 |
| Evolution | Vol. 72 (3); 2018 | Vol. 70 (12); 2016 |
| Mol. Biol. Evol. | Vol. 35 (3); 2018 | Vol. 33 (6); 2016 |
| Mol. Phylogenet. Evol | Vol. 120; 2018 | Vol. 93; 2015 |
| New Phytol. | Vol. 217(3); 2018 | Vol. 217(1); 2018 |
| Syst. Biol. | Vol. 67 (1); 2018 | Vol. 64 (1); 2015 |
| Syst. Bot. | Vol. 42(4); 2017 | Vol. 42(1); 2017 |

